# Cellulosic copper nanoparticles and a hydrogen peroxide-based disinfectant protect Vero E6 cells against infection by viral pseudotyped particles expressing SARS-CoV-2, SARS-CoV or MERS-CoV Spike protein

**DOI:** 10.1101/2022.03.22.485373

**Authors:** Ariane Brault, Raphael Néré, Jérôme Prados, Simon Boudreault, Martin Bisaillon, Patrick Marchand, Patrick Couture, Simon Labbé

**Affiliations:** Département de Biochimie et de Génomique Fonctionnelle, Faculté de médecine et des sciences de la santé, Université de Sherbrooke, Sherbrooke, QC, J1E 4K8, Canada; SaniMarc Group, Victoriaville, QC, G6P 7E3, Canada

**Keywords:** Murine leukemia virus, coronavirus, pseudoviral particles, envelope glycoprotein Spike, copper nanoparticles, disinfectant.

## Abstract

Severe acute respiratory syndrome (SARS) is a viral respiratory infection caused by human coronaviruses (HuCoV) that include SARS-CoV-2, SARS-CoV, and Middle East respiratory syndrome coronavirus (MERS-CoV). Although their primary mode of transmission is through contaminated respiratory droplets from infected carriers, the deposition of expelled virus particles onto surface and fomites could contribute to viral transmission. Here, we use replication-deficient murine leukemia virus (MLV) pseudoviral particles expressing SARS-CoV-2, SARS-CoV, or MERS-CoV Spike (S) protein on their surface. These surrogates of native coronavirus counterparts serve as a model to analyze the S-mediated entry into target cells. Carboxymethyl cellulose (CMC) nanofibers that are combined with copper (Cu) exhibit strong antimicrobial properties. S-pseudovirions that are exposed to CMC-Cu nanoparticles (30 s) display a dramatic reduction in their ability to infect target Vero E6 cells, with ∼97% less infectivity as compared to untreated pseudovirions. In contrast, addition of the Cu chelator tetrathiomolybdate protects S- pseudovirions from CMC-Cu-mediated inactivation. When S-pseudovirions were treated with a hydrogen peroxide-based disinfectant (denoted Saber^TM^) used at 1:16 dilution, their infectivity was dramatically reduced by ∼98%. However, the combined use of Saber^TM^ and CMC-Cu is the most effective approach to restrict infectivity of SARS-CoV-2-S, SARS-CoV-S, and MERS-CoV-S pseudovirions in Vero E6 cell assays. Together, these results show that cellulosic Cu nanoparticles enhance the effectiveness of diluted Saber^TM^ sanitizer, setting up an improved strategy to lower the risk of surface- and fomite-mediated transmission of enveloped respiratory viruses.

## Introduction

The emergence of the novel human severe acute respiratory syndrome coronavirus 2 (SARS-CoV-2) represents a severe public health burden worldwide.^1–4^ The fast spread of SARS-CoV-2 pandemic poses a striking challenge for public services and health facilities. Public spaces and healthcare environments become rapidly contaminated by way of person-to-person contact. Several studies have shown that the main way by which people are infected with SARS-CoV-2 is through respiratory droplet transmission and airborne spread.^5–9^ Individuals can also be infected with SARS-CoV-2 through contact with surfaces and inanimate materials, called fomites. However, the risk of surface-mediated transmission is dependent on several factors.^10–15^ Among these are the infection prevalence rate in the community, the density of virus particles present on the surfaces, the time between viral contamination of the surface and transmission to the people, and the amount of virus particles from surfaces to hands and from hands to the mucous membranes of the nose, mouth, and eyes.

Coronaviruses including SARS-CoV-2 remain viable on a variety of porous and non-porous surfaces.^10, 16–18^ Although these viruses are rapidly eliminated on porous surfaces within minutes to few hours, they are more persistent on non-porous surfaces such as plastic, glass, and stainless steel in which case they can remain infectious up to 3 days.^10, 17, 19, 20^ In contrast, SARS-CoV-2 is rapidly inactivated after 4 h on copper (Cu) surfaces.^18^ Interestingly, yeasts, bacteria, and viruses are also rapidly killed on surfaces of Cu or Cu alloys containing at least 70% Cu.^21–23^ This process is called Cu-mediated “contact killing”.^21^ Importantly, it has also been known that concentrations of Cu required to kill microbes are not toxic to humans.^24^ The insensitivity of human tissue to Cu can be contrasted with microorganisms that are extremely sensitive to its toxic effects.^25^

One exciting new application of nanotechnology consists of using carboxymethyl cellulose (CMC) as a nanosubstrate to generate Cu-containing nanoparticles using copper sulfate and sodium borohydride.^26, 27^ The resulting cellulosic cuprous nanoparticles (CMC-Cu) possess antimicrobial activity against non-pathogenic microbes such as the bacterium *Escherichia coli* and the yeast *Saccharomyces cerevisiae*.^26, 28, 29^ Biophysical analysis of CMC-Cu nanoparticles have revealed that Cu is incorporated into the CMC polymer in its reduced state (Cu^1+^).^27^ Electron microscopic analyses have shown the presence of spherical particles on the surface of the CMC film.^26^ Cu is thought to be slowly released from these particles under its reduced state (Cu^1+^) and becomes highly toxic as it changes its oxidative state. This redox active process is predicted to be associated with generation of destructive hydroxyl radical and superoxide anions. These reactive oxygen species (ROS) have the potential to cause detrimental oxidative damage to vital microbial cell constituents such as lipids (including those in the cell membrane), proteins and nucleic acids. For example, in the case of *S. cerevisiae*, results have shown that it is more sensitive to CMC-Cu nanoparticles than soluble CuSO_4_.^28^ Yeast cells treated with CMC-Cu are more susceptible to lipid peroxidation than untreated cells. Furthermore, the CMC-Cu-treated cells exhibit morphological anomalies that are indicative of cell surface damage and loss of membrane integrity.^29^ These studies suggest the potency of CMC-Cu as sanitizing agent to prevent contamination of fomites by enveloped coronaviruses is of interest.

An efficient way to prevent indirect community transmission of coronaviruses by contaminated surfaces consists of using disinfectants and cleaning agents against enveloped viruses.^30^ World Health Organization (WHO) recommends that disinfectants must contain 70% ethanol or isopropanol, or 0.5% hydrogen peroxide and, must be used at least, a 20 s treatment.^31, 32^ One caveat of using alcohol-based disinfectants is their high volatility.^33^ To favor the time of retention of alcohol-based disinfectants on inanimate surfaces, anionic surfactants such as sodium dodecylbenzenesulfonate and sodium laureth sulfate must be added.^33^ Moreover, owing to their intrinsic nature that includes both hydrophobic and hydrophilic properties, anionic surfactants possess the advantage that they can dissolve the outer lipid layer of enveloped viruses, while remaining soluble in water as well as in alcohol- and H_2_O_2_-based disinfectants.^34, 35^

Studies using native coronaviruses such as SARS-CoV-2, SARS-CoV, and MERS-CoV are restricted to biosafety level 3 (BSL-3) containment facilities.^36^ However, alternative models have been developed that allow production of replication-deficient enveloped virus-like particles that are safer surrogates of native coronaviruses.^36–39^ One useful and safe model is to use murine leukemia virus (MLV) core components to form the enveloped pseudovirion backbone.^36, 38^ During its formation, for instance in HEK-293T cells, a second expression plasmid triggers the synthesis of a luciferase reporter transcript flanked by retroviral regulatory LTR regions and a packaging signal that allow incorporation of the reporter mRNA into a nascent virus-like particle. Moreover, during the assembly process a third plasmid encodes either SARS-CoV-2, SARS-CoV, or MERS-CoV Spike (S), resulting in MLV-based pseudoviral particles harboring coronavirus S proteins on their surface. These pseudovirions are competent to infect susceptible cells and their entry is specifically dictated by the SARS-CoV-2-, SARS-CoV- or MERS-CoV-S protein present on their surface. These S-pseudovirions are useful surrogates in studies focusing on entry events of enveloped viruses. Following entry and infection of target cells, the luciferase reporter becomes integrated in the host cell genome and is expressed, thus enabling quantification of virus entry events of S-pseudotyped particles.

In this report, SARS-CoV-2-, SARS-CoV- and MERS-CoV-S-pseudotyped particles were rapidly inactivated upon exposure to CMC-Cu. That resulted in a dramatic decrease of their potency to infect target cells. Similarly, infectivity of these three types of S-pseudotyped particles was dramatically reduced when they had been pretreated with the disinfectant Saber^TM^ over a 30 s time period. Of significance, results showed that a pretreatment with Saber^TM^ in combination with CMC-Cu was the most effective strategy to prevent infectivity of SARS-CoV-2-, SARS- CoV- and MERS-CoV-S pseudovirions as well as pseudoviral particles expressing a relevant variant form of SARS-CoV-2 S protein.

## Materials and methods

### Cell culture

HEK-293T (human) and Vero E6 (African green monkey) cells were grown in Dulbecco’s modified Eagle’s medium (DMEM; Wisent). DMEM medium was supplemented with 10% fetal bovine serum (FBS; Wisent) and 1X penicillin-streptomycin (PS) solution (Mediatech). Cell lines were grown at 37°C in the presence of 5% CO_2_ under a humidified atmosphere. Transfection of HEK-293T cells was carried out using Lipofectamine 2000 (Thermo Fisher Scientific - Invitrogen). Transfection conditions used a ratio of 1 µg total DNA (including all three plasmids) for 3 µl Lipofectamine 2000 that were mixed in the presence of Opti-MEM (Life Technologies) as described previously.^36, 38^ Cell viability was determined by a colorimetric MTT (3-(4,5-dimethylthiazol-2-yl)-2,5-diphenyltetrazolium bromide) assay as described previously.^40, 41^ Untreated cells were used as controls and these samples were defined as 100% cell viability. The 50% inhibitory concentrations (IC_50_) were determined graphically using GraphPad Prism 9.3.1 software in which calculations were made by considering concentration of Saber^TM^ on the X axis on a logarithmic scale and relative cell viability on the Y axis, on a linear scale. All assays were performed in triplicate and repeated at least three times from independent experiments.

### DNA constructs

Plasmid pcDNA3.1-SARS-CoV-S contained a short coding sequence corresponding to the CD5 signal peptide (codons 1-24) that replaced the first eleven N-terminal codons in the original SARS-CoV-S sequence. To incorporate the CD5 signal peptide DNA fragment in-frame with the coding sequences of MERS-CoV-S and SARS-CoV-2-S-Δ19, we performed PCR amplification of the DNA fragment corresponding to the CD5 signal peptide found in pcDNA3.1-SARS-CoV-S using primers designed to generate HindIII and EcoRI or NotI and ApaI restriction sites at the upstream and downstream termini of the DNA fragment. The resulting PCR products were digested with HindIII and EcoRI or NotI and ApaI, and were then cloned into the HindIII-EcoRI-digested pcDNA3.1-MERS-CoV-S or NotI-ApaI-digested pcDNA3.1-SARS-CoV-2-S-Δ19 vectors. The resulting plasmids were named pcDNA3.1-CD5-MERS-CoV-S and pcDNA3.1-CD5-SARS-CoV-2-S-Δ19, respectively. In the case of pcDNA3.1-CD5-MERS-CoV- S, the DNA fragment insertion resulted in the replacement of the first fifteen codons of MERS-CoV-S by the CD5 signal (codons 1-24). Similarly, the first eleven codons of SARS-CoV-2-S- Δ19 were replaced by the CD5 signal in the pcDNA3.1-CD5-SARS-CoV-2-S-Δ19 plasmid. SARS-CoV-2-S mutant alleles were created to substitute the codons corresponding to Asn501 and Asp614 with Tyr and Gly codons, respectively. These N501Y and D614G mutants were generated by a PCR overlap extension method.^42^ DNA sequences containing site-specific mutations from each respective PCR-amplified fragment was digested with NotI and ApaI and exchanged with a corresponding DNA region into the pcDNA3.1-CD5-SARS-CoV-2-S-Δ19.

### Synthesis of Luciferase (LUC) RNA molecules for production of standard curves

Using plasmid pTG-Luc as a template, the DNA sequence encoding *LUC* was isolated by PCR with primers containing KpnI and BamHI sites.^36^ The PCR product was cloned into the corresponding sites of pBluescript SK in which a T7 promoter sequence is found immediately next to the cloning site on the 5’ end. The resulting plasmid was named pSK-Luc. Using BamHI-linearized pSK-Luc (5 µg per reaction), in vitro run-off transcription with T7 RNA polymerase was performed using buffer T (80 mM HEPES, pH 7.5, 20 mM MgCl_2_, 2 mM spermidine, 40 mM DTT and 10 mM NaCl) that was supplemented with pyrophosphatase (0.1 U/µl) as described previously.^43^ The DNA template was eliminated by digestion with RNase-free DNase I for 15 min at 37°C. *LUC* RNA molecules were purified using preparative polyacrylamide gel electrophoresis as described previously ^44, 45^. Subsequently, RNA molecules were extracted from the gel using the crush and soak method.^45^ Elution of RNA was performed in buffer E (2 M ammonium acetate, 1% sodium dodecyl sulphate and 1 µg glycogen) for 18 hr at 55°C. After a centrifugation to remove gel debris, the supernatant containing eluted RNA was collected and precipitated with ethanol. After collecting the RNA by centrifugation, it was quantified spectrophotometrically at 260 nm. After RNA levels quantification, measurements of transcripts were converted to the molecule number per μl as described previously.^46^ Serial 10-fold dilutions of the quantified *LUC* RNA (calculated in RNA transcript copies per reaction) were prepared and used as standards for the generation of standard curves to determine the limit of detection (LoD) of the primer set in RT-qPCR assays.

### Quantification of transcript levels by RT-qPCR

Total RNA was extracted from freshly produced S-pseudoviral particles or HEK-293T cells using a TRIzol method.^47^ Reverse transcription reactions were performed in a 20-µl reaction mixture that contained the specified amount of RNA transcripts produced in vitro or 1 µg of RNA extracted from either HEK-293T cells or S-pseudovirus preparations, 2 µl of random primer mix (60 µM; 25 µM oligo(dT) and 35 µM random hexamers), 2 µl 10X RT buffer, 1 µl dNTP mix (10mM) (Thermo Fisher - Invitrogen), 0.2 µl RNase inhibitor (40 U/µl), and 0.2 µl MMuLV RT (200 U/µl). cDNA synthesis was performed using the following steps: 5 min at 25°C for hybridization, 60 min at 42°C for elongation of cDNA, and 20 min at 65°C for inactivation. qPCR reactions were performed in a 20-µl reaction mix that contained 2 µl of cDNA (1:10 dilution), 300 nM of forward and reverse specific primers, and 10 µl of Supermix qPCR 2X that included SYBR Green, dNTPs and the thermostable DNA polymerase. The reaction was performed on a CFX96 Touch Real-Time PCR System (Bio-Rad) using the following steps: 3 min at 95°C for the initial denaturation, 15 s at 95°C for denaturation, 30 s at 60°C for hybridization, and 30 s at 72°C for elongation. The last three steps were repeated over 45 cycles. Fluorescence measurements were monitored after each amplification round and the threshold cycle (C_t_) value for each sample was calculated by assessing the point at which fluorescence crossed the threshold line, exhibiting an increase in fluorescence above the calculated background levels. The result was considered valid if the target-specific fluorescent signal showed the C_t_ value ≤ 37 cycles and all positive and negative control reactions gave a successful and no amplification, respectively. MIQE guidelines were followed to perform and interpret RT-qPCR results.

### Production of S-pseudoviral particles

The biological system to produce S-pseudotyped viral particles was based on a replication-deficient murine leukemia virus (MLV) that lacked essential genes to achieve a complete replication cycle.^36, 38^ Furthermore, the genome of viral particles was modified to encode the *LUC* reporter gene, thereby allowing measurement of infectivity of the S-pseudoviral particles after host cell infection. For production of MLV-based pseudoviral particles, HEK-293T cells were co-transfected with three plasmids as described previously.^36, 38^ One plasmid encoded the MLV core genes *gag* and *pol* but lacked the MLV envelope *env* gene. A second plasmid harbored the *LUC* reporter gene and packaging viral sequences that allowed this template to be encapsulated into nascent pseudoviral particles. A third plasmid encoded the desired envelope glycoprotein such as SARS-CoV-S, SARS-CoV-2-S, MERS-CoV-S, and VSV-G (positive control), or was an empty expression vector (negative control). Co-expression of these three plasmids in HEK-293T allowed production of pseudoviral particles expressing the indicated coronavirus S protein under conditions that were described previously.^36, 38^ The decanoyl-RVKR- CMK inhibitor (37.5 µM) of proprotein convertases was added to HEK-293T cells immediately following co-transfection of plasmids and boosted with an additional aliquot of inhibitor (37.5 µM) 24 h after transfection.^48^ Two days following co-transfection, cells and cell debris were harvested and supernatants containing S-pseudoviral particles were filtered and either immediately used for infection of VeroE6 cells or aliquoted and stored at -80°C for further infections of target cells.

### Protein analysis of pseudoviral particles

In the case of detection of VSV-G and coronavirus S proteins on the surface of pseudoviral particles and the capsid p30 protein of MLV, freshly produced pseudotyped viral particles were filtered and divided into 3 ml aliquots. Pseudoviral particles were purified from these aliquots by ultracentrifugation at 50,000 rpm for 2 h. The pelleted virus-like particles were resuspended, dissolved in sample buffer (120 mM Tris-HCl, pH 6.8, 20% glycerol, 4% sodium dodecyl sulfate (SDS), 1.4 mM β-mercaptoethanol, 500 µM dithiothreitol (DTT), 3 M urea, and 1 M thiourea) and heated at 65°C for 20 min prior to their protein content separation on 4 – 6% or 4 – 12% gradient SDS-polyacrylamide gels. After electrophoresis, samples were analyzed by immunoblot assays using the following primary antibodies: monoclonal anti-p30 antibody Ab130757 (Abcam); monoclonal anti-C9 antibody Ab5417 (Abcam) (used for C9-tagged S proteins of SARS-CoV and MERS-CoV); polyclonal anti-VSV-G antibody EPR12997 (Abcam) and polyclonal anti-SARS-CoV-2 S antibody 40590-T62 (SinoBiological). Following incubation with primary antibodies, membranes were washed and incubated with the appropriate horseradish peroxidase-conjugated secondary antibodies (Amersham Biosciences), developed with enhanced chemiluminescence reagents and visualized by chemiluminescence using an ImageQuant LAS 4000 instrument (GE Healthcare) equipped with a Fujifilm High Sensitivity F0.85 43 mm camera.

### Infection of Vero E6 cells by pseudoviral particles and luciferase assays

Vero E6 cells were cultured and prepared to seed cells to wells of a 24-well plate by adding 2.5 X 10^5^ cells in complete DMEM as described previously.^36, 38^ After 18 h, cells were washed three times with 0.5 ml pre-warmed Dulbecco’s PBS (DPBS) medium and then infected with pseudoviral particles (200 µl) harboring the desired S protein as described previously.^36, 38^ Cells were incubated on a rocker for 2 h and then supplemented with complete DMEM (300 µl) medium and incubated for 72 h. Infected cells were lysed as described previously.^36, 38^ Lysed cell supernatant from each well was assessed for luciferase activity using the Luciferase Assay System (Promega). Relative luciferase units (RLU) were determined using the Glomax 20/20 luminometer (Promega) and infectivity values were analyzed as described previously ^36^ and plotted using GraphPad Prism 9.3.1 software.

### Preparation of carboxymethyl cellulose copper (CMC-Cu) nanoparticles

Sodium (Na)–carboxymethyl cellulose (CMC) (0.2 g) was dissolved in 9.8 ml of sterile MilliQ water under vigorous magnetic stirring conditions as described previously.^26^ Three milliliters of CuSO_4_ (0.1 M) were added by dropwise into the Na-CMC solution under continuous stirring. CuSO_4_ was added to final concentration of 20 mM. The mixture was left at room temperature for 14 h without stirring. The following step consisted in reducing Cu^2+^ ions of the CMC-Cu composite material by slowly adding 2 ml of sodium borohydride (NaBH_4_, 0.5 M solution in diglycine) over a span of 30 min. The resulting cellulose copper nanoparticles were used immediately at the appropriate dilution.

### Treatments of pseudoviral particles

Exposure of pseudoviral particles to Saber^TM^ was performed using freshly produced MLV-based pseudotyped particles (500 µl) that were incubated with Saber^TM^, which was diluted at ratios of 1:250 and 1:100. Pseudoviral particles were exposed to the disinfectant for 30 s or 15 min. Following the exposure times, samples were gel-filtered twice through a Sephadex LH-20 column and the filtrate containing pseudoviral particles (200 µl) was inoculated onto Vero E6 cells. After incubation on a rocker for 2 h, cells were supplemented with complete DMEM (300 µl) and incubated for 72 h. Infected cells were lysed as described previously.^36, 38^ Lysed cell supernatant (10 µl) from each well was assessed for luciferase activity using the Luciferase Assay System.

CMC-Cu nanoparticles containing either 200 µM, 400 µM or 800 µM copper were used to treat MLV-based pseudotyped particles (1 ml) for 30 s or 15 min. Samples were then filtered through a 0.22 µM pore-sized membrane to remove CMC-Cu nanoparticles. The filtrate containing pseudoviral particles (200 µl) was inoculated onto Vero E6 cells and the cells were then incubated on a rocker for 2 h prior to addition of complete DMEM (300 µl). Vero E6 cells were then incubated for 72 h and luciferase assays performed as described previously.^36, 38^ CMC-Cu nanoparticles were preincubated (5 min) with the copper chelator TTM (4 mM) or the antioxidant compound Tiron (100 mM) when indicated. To analyze data, the luciferase activity of infected cells with SARS-CoV-2- or SARS-CoV- or MERS-CoV-S pseudovirions without CMC-Cu treatment was set up to 100% and luciferase activities of cells infected with CMC-Cu-treated S- pseudovirions were measured and compared to untreated S-pseudovirions.

## Results

### Infection of Vero E6 cells with pseudoviral particles expressing the S protein of SARS-CoV-2, SARS-CoV, and MERS-CoV

During the production of MLV-based virus-like particles, we assumed that the RNA molecule being encapsulated into pseudoviral particles contained only one copy of the coding sequence of the luciferase (*LUC*) gene reporter. Based on this assumption, we isolated the coding region of *LUC* and subcloned it in a way that a T7 promoter sequence was located at its 5’ end. The resulting DNA template was used to generate synthetic *LUC* transcripts that were utilized as standards in RT-qPCR analysis to produce standard curves by linear regression (Fig. 1A). This approach allowed us to evaluate the number of *LUC* RNA transcripts that were present in any given batch of pseudoviral particles. This in turn allowed calibration and comparison of infectivity of pseudotyped virions with various types of viral envelope glycoproteins.

**Fig. 1.**
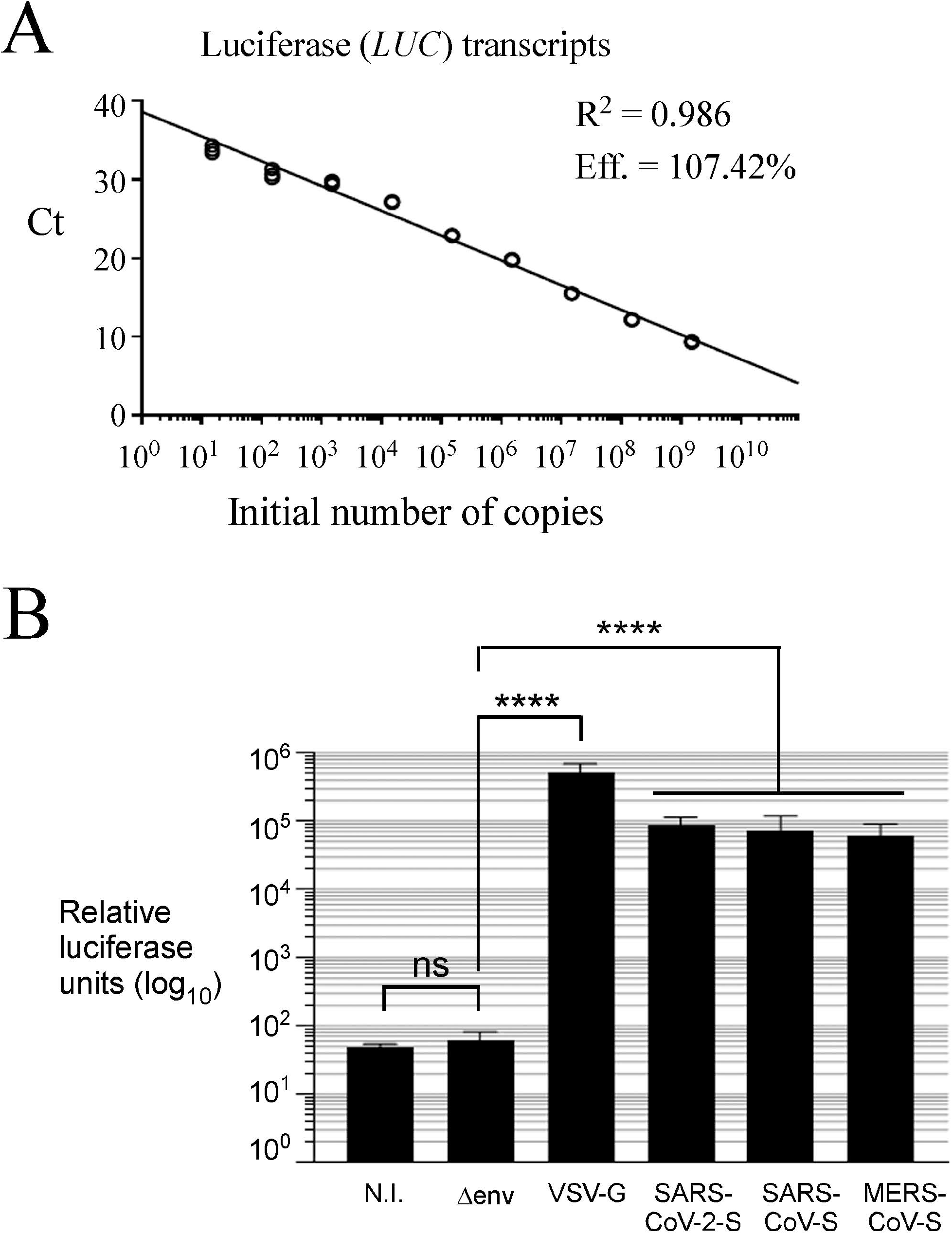
Infectivity assays using a MLV-based system that produces pseudoviral particles expressing the indicated coronavirus S protein in Vero E6 cells. *A*, Representative linear regression curve to assess *LUC* RNA copy number in infectivity assays. Tenfold serial dilutions from 10^1^ to 10^9^ copies of *LUC* synthetic RNA transcripts were analyzed by RT-qPCR assays. The graph was generated by plotting the C_t_ values (y-axis) and the logarithm of initial number of copies of synthetic transcripts (x-axis). Calculated linear correlation coefficients (R^2^) and percentage of amplification efficiencies (Eff) are indicated for the primer set. The graph shows data for three independent experiments. The different values obtained for each point were very similar and are superimposed for most data points. *B*, Quantification of luciferase activity of lysed Vero E6 cells 72 h post-infection. Pseudoviral particle codes are as follows: N.I., non-infected control; Δenv, pseudoviral particles lacking a viral envelope glycoprotein; VSV G, VSV G-pseudotyped particles; SARS-CoV-2-S, SARS-CoV-2-Spike-pseudotyped particles; SARS-CoV-S, SARS- CoV-Spike-pseudotyped particles; MERS-CoV-S, MERS-CoV-Spike-pseudotyped particles. Results show the average relative luciferase units (log_10_) of a minimum of five independent experiments. Error bars indicate standard deviation (± SD; error bars). The asterisks correspond to *p* < 0.0001 (****) (one-way ANOVA with Dunnett’s multiple comparisons test against the Δenv control), whereas ns stands for not significant.

Experimental setup using equal amounts of pseudoviral particles determined that 1.0 x 10^9^ viral copies were used per viral infection. In the case of VSV-G-pseudotyped particles that were used as positive control particles, results showed that their viral infection of target cells produced 5.2 x 10^5^ relative luciferase units (RLU) (Fig. 1B). In contrast, non-infected (N.I.) cells or cells infected with Δenv pseudoviral particles lacking a cell-surface viral envelope glycoprotein (negative control) gave low background levels of RLU (4.9 x 10^1^ RLU for N.I. cells and 6.3 x 10^1^ RLU for Δenv particles) (Fig. 1B). In the case of SARS-CoV-2-, SARS-CoV-, and MERS-CoV-S-pseudotyped particles used to infect Vero E6 cells, luciferase assays (which serve to quantify the infectivity of pseudoviral particles) reached 8.7 x 10^4^, 7.6 x 10^4^, and 6.1 x 10^4^ RLU, respectively. These results were significantly higher compared to baselines of RLU in N.I. cells and cells infected with Δenv particles (Fig. 1B). Indeed, these results showed a ≥3-logarithmic increase in the case of SARS-CoV-2-, SARS-CoV-, and MERS-CoV-S-pseudotyped particle infectivity in comparison with background levels of infectivity in the case of Δenv pseudoviral particles in Vero E6 cells.

### Detection of the presence of coronavirus S mRNA and proteins during generation of pseudotyped particles

We analyzed the profiles of expression of the S transcript and its corresponding S protein as well as the VSV-G mRNA and its protein product (control) during production of S-pseudoviral particles in HEK-293T cells. The levels of VSV-G and S mRNAs were analyzed (RT-qPCR assays) 24 h and 48 h after co-transfection of plasmids in producer HEK-293T cells. Results showed that VSV-G and S transcripts were expressed at both times (Fig. 2A). Although S transcript levels exhibited a slight reduction at 48 h compared to 24 h, their mRNA levels remained robustly expressed. In the case of VSV-G transcripts that were detected at 48 h, results showed that their levels were reduced 3.7-fold less than those observed at 24 h (Fig. 2A). However, this decrease was not sufficient to prevent detection at 48 h of both VSV-G mRNA and protein steady-state levels (Fig. 2B).

**Fig. 2.**
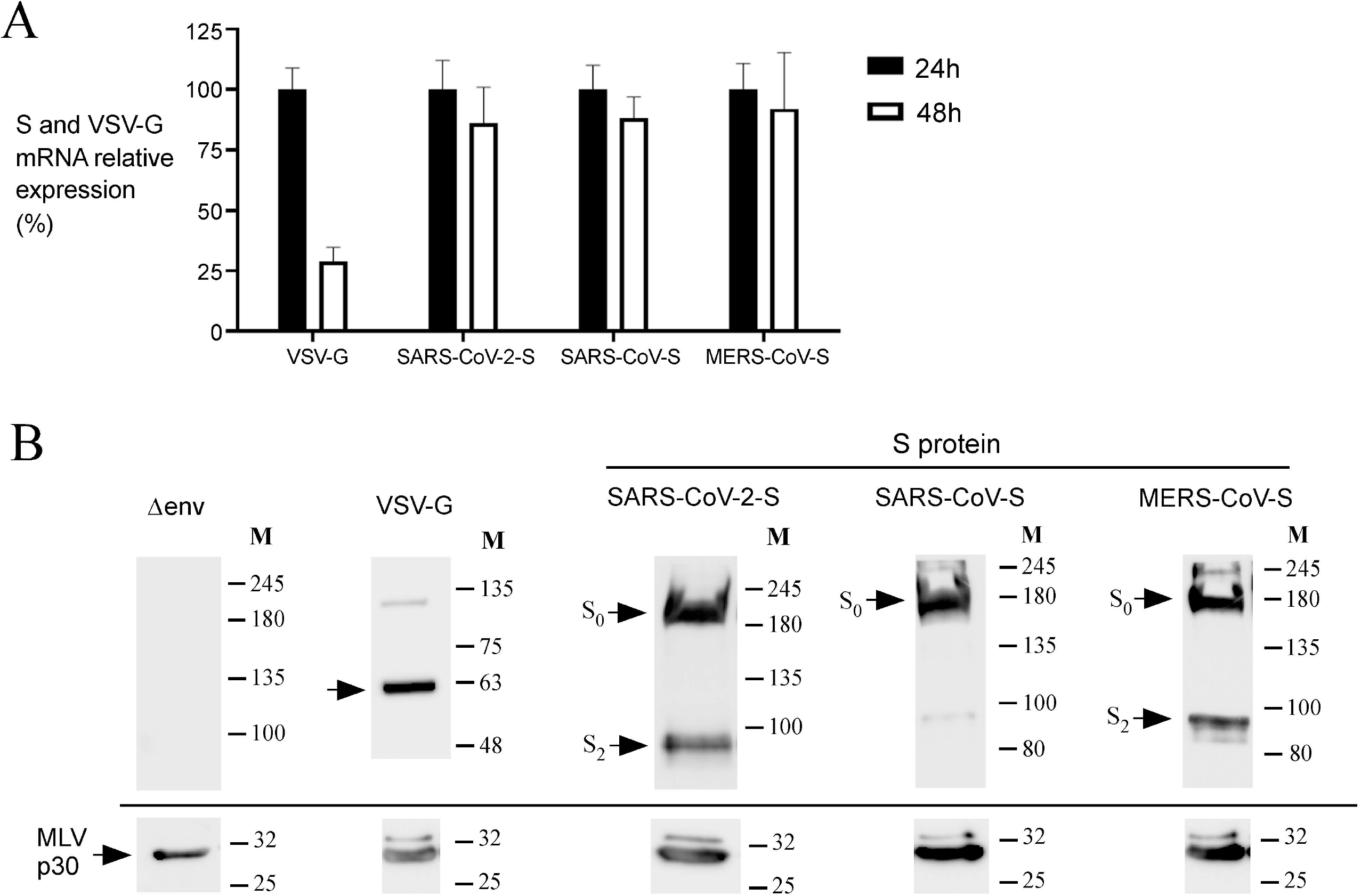
Assessment of S expression in a MLV-based coronavirus pseudotyped particle production system. *A*, Representative RT-qPCR analysis of RNA samples isolated following transfection of plasmids expressing *S* genes of SARS-CoV-2, SARS-CoV, and MERS-CoV, or VSV-G gene of the rhabdovirus vesicular stomatitis virus (VSV). RNA samples were isolated from transfected HEK-293T cells after 24 h and 48 h. Specific primer sets targeting unique regions of VSV-G, SARS-CoV-2-S, SARS-CoV-S, and MERS-CoV-S genes were used to probe their corresponding transcripts. A specific primer set was used to detect human β_2_-microglobulin (B2M) transcript as an internal control for normalization during quantification of the RT-qPCR products. *B*, Western blot analysis of the S protein of SARS-CoV-2, SARS-CoV, and MERS-CoV as well as the VSV- G and MLV p30 (control) proteins after 48 h transfection. S_0_, uncleaved form of S protein; S_2_, S2 segment of S protein. The positions of the molecular weight standards (in kDa) are indicated on the right side.

S-pseudoviral particles were retrieved after 48 h of transfection to further validate the presence of S proteins of SARS-CoV-2, SARS-CoV, and MERS-CoV. Purified virus-like particles were analyzed by immunoblot assays. Results showed that S proteins were present in SARS-CoV-2, SARS-CoV, and MERS-CoV pseudovirions (Fig. 2B). In the case of SARS-CoV-2-S and MERS-CoV-S, the uncleaved (S_0_) form and the band corresponding to the S2 segment (S_2_) were detected as previously reported (Fig. 2B).^48, 49^ The presence of the lower molecular weight band S_2_ could have resulted from the action of furin or some related proprotein convertase(s) activity during the process of maturation that occurs within the secretory pathway in HEK-293T cells. Weakly detected higher order bands may correspond to oligomeric forms of S protein.^48, 49^ VSV-G and S proteins were undetectable in Δenv pseudoviral particles that are devoid of viral envelope glycoproteins (Fig. 2B). Production of the p30 capsid protein of MLV was co-currently followed as a control for each pseudotyped particle preparation. This approach was used to validate the presence of an additional viral protein produced and incorporated into MLV-based pseudoviral particles. Taken together, these results indicated that pseudoviral particles contain the viral envelope VSV-G or S protein that is required to infect Vero E6 cells.

### Inhibition of SARS-CoV-2-, SARS-CoV-, and MERS-CoV-S-pseudotyped particle infection by Saber^TM^

MLV-based pseudoviral particles harboring the S protein of SARS-CoV-2, SARS-CoV and MERS-CoV were associated with increased viral entry in Vero E6 cells highlighted by luciferase activity. We assessed whether pretreating S protein-bearing pseudoviral particles with the disinfectant Saber^TM^ decreased the sensitivity of Vero E6 cells to pseudoviral particles. Prior to these experiments, the cytotoxic effect of Saber^TM^ alone (without the presence of pseudoviral particles) on Vero E6 cells was determined in this manner. Vero E6 cells were treated with 0.1, 0.2, 0.4, 1.0, 2.0, 2.5, 4.0, and 5.0% of Saber^TM^ for 30 s. After a gel filtration step followed by 72 h of incubation, cell viability was assessed by a MTT method (Fig. S1, A).^41^ The calculated concentration of Saber^TM^ that resulted in half maximal inhibitory concentration (IC_50_) was 1.48%, whereas 1.0% and lower concentration of Saber^TM^ did not significantly affect Vero E6 cell viability (Fig. S1, A).

We next tested the effect of Saber^TM^ on the purified pseudoviral particles expressing different versions of S protein prior their assays to infect target cells. In the case of SARS-CoV-2-S-pseudotyped particles, they were treated with 0.0004, 0.0005, 0.001, 0.0013, 0.002, 0.004, 0.01, 0.02, 0.04, 0.07, 0.1, 0.2, 0.4 and 1.0% of Saber^TM^ over a period of time of 30 s. In the case of SARS-CoV-S-pseudovirions, Saber^TM^ concentrations varied from 0.0006 to 2.0% for the same time of treatment. In the case of MERS-CoV-S-pseudovirions, these were treated with Saber^TM^ in concentration ranging from 0.001 to 1%. Saber^TM^-treated S-pseudovirions were used to infect Vero E6 cells, and a quantification of their cellular entry by luciferase activity assays. Results showed that the Saber^TM^ concentration required for a 50% inhibition (IC_50_) of infectivity of viral SARS-CoV-2-, SARS-CoV- and MERS-CoV-S-pseudotyped particles was 0.02%, 0.06%, and 0.04%, respectively (Fig. 3, A – C). Together, these results revealed that SARS-CoV-2-, SARS- CoV- and MERS-CoV-S-pseudotyped particles were 74-, 25-, and 37-fold more sensitive to Saber^TM^ than Vero E6 cells were.

**Fig. 3.**
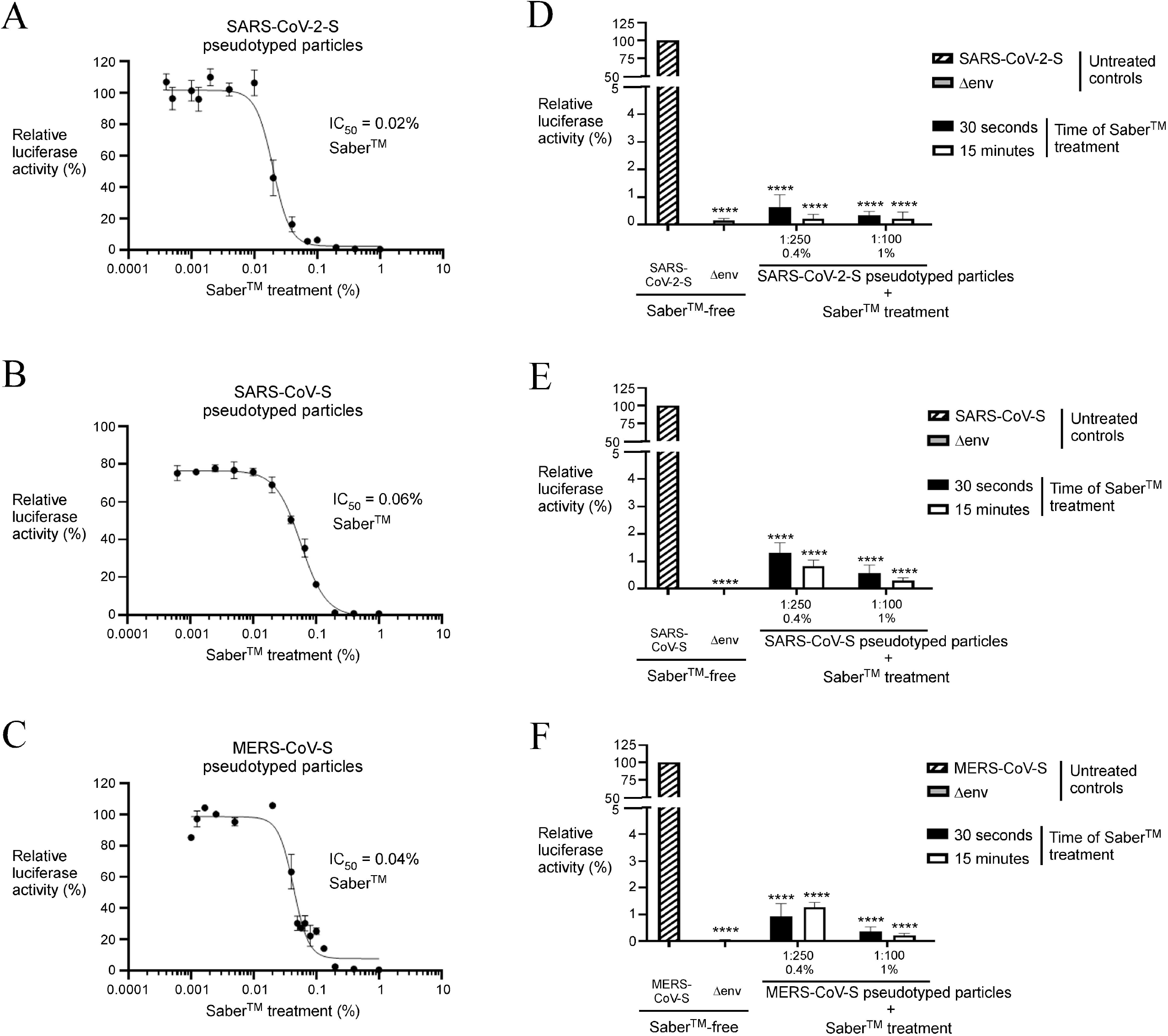
Saber^TM^ protects Vero E6 cells against infectivity of SARS-CoV-2-, SARS-CoV-, and MERS-CoV-S-pseudotyped virions. *A – C*, SARS-CoV-2-, SARS-CoV-, and MERS-CoV-S- pseudotyped particles were treated with different doses (ranging from 0.0004 to 1%) of Saber^TM^. After a 30 s treatment, S-pseudovirions were used to infect Vero E6 cells. Quantification of the infectivity of Saber^TM^-treated S-pseudovirions in infected cells was measured by luciferase assays. *D – F*, The S-pseudotyped virions were incubated in the presence of Saber^TM^ that was prepared in a specific volume of a dilute solution 1:250 (0.4%) and 1:100 (1%) from a concentrated stock solution. S-pseudovirions treated with Saber^TM^ over a period of 30 s or 15 min were used to infect Vero E6 cells. As controls, Vero E6 cells were infected with untreated S-pseudoviral particles (Saber^TM^-free, positive control) or pseudovirions lacking a cell-surface viral envelope glycoprotein (Δenv, negative control). Luciferase activity was measured 72 h post infection. Luciferase signal of infected cells with untreated S-pseudovirions (Saber^TM^-free) was set up to 100%. Graphs represent quantification of the results of at least five independent infection experiments performed in biological triplicate. Values are represented as averages ± S.D. The asterisks correspond to *p* < 0.0001 (****) (one-way ANOVA with Dunnett’s multiple comparisons test against the Saber^TM^- free positive control).

We therefore used concentrations of 0.4 and 1.0% Saber^TM^ to pretreat S-pseudoviral particles prior to infect Vero E6 cells. These Saber^TM^ concentrations were largely in excess to inactivate pseudovirions but not sufficiently high to affect Vero E6 cell viability. Results showed that Saber^TM^ strongly decreased infectivity of SARS-CoV-2-, SARS-CoV- and MERS-CoV-S-pseudotyped particles after 30 s and 15 min of treatment (Fig. 3, D – F). The inhibitory effect of Saber^TM^ on the cellular entry of S-pseudovirions was already obvious even at the lowest dilution (1:250, 0.4% Saber^TM^) used in the assays (Fig 3, D – F). When Vero E6 cells had been infected with Saber-treated pseudoviral particles expressing SARS-CoV-2 S protein, results showed significant decreases of 99.4% (30-s exposure) and 99.8% (15-min exposure) of luciferase activity compared to control conditions (untreated pseudoviral particles) (Fig. 3D). The inhibitory effect of Saber^TM^ to restrain cellular entry of SARS-CoV-2-S-pseudotyped virions was maintained when it was used at a 1% (1:100 dilution) over periods of time of 30 s and 15 min. Under these conditions, lysates of Vero E6 cells exhibited 99.6% and 99.8% less luciferase activity as compared to the Saber^TM^-free control condition (Fig. 3D).

Similarly, in the case of SARS-CoV-S pseudotyped particles, a pretreatment with 0.4% or 1% Saber^TM^ for 30 s or 15 min strongly inhibited their ability to infect Vero E6 cells (Fig. 3E). Results showed that cell extracts from Vero E6 cells that had been exposed to Saber^TM^-treated SARS-CoV-S pseudoviral particles exhibited 98.7% to 99.7% less luciferase activity than conditions without Saber^TM^ treatment (Fig. 3E). MERS-CoV-S pseudoviral particles also exhibited hypersensitivity to 0.4% and 1% concentrations of Saber^TM^ treatment for 30 s or 15 min (Fig. 3F). Utilization of Saber^TM^-treated MERS-CoV-S pseudoviral particles dramatically lowered the relative luciferase activity that was measured in cell extracts prepared from infected Vero E6 cells. Saber^TM^ concentration as low as 0.4% inhibited luciferase activity in target Vero E6 cells by 98.7% (Fig. 3F), whereas luciferase activity was reduced by 99.6% in the presence of 1% Saber^TM^ (Fig. 3F). Taken together, these results indicated that Saber^TM^ is an efficient disinfectant to reduce SARS-CoV-2-, SARS-CoV-, and MERS-CoV-S-pseudotyped particle infection in a MLV-based pseudoviral system.

### CMC-Cu nanoparticles inhibit viral infection of Vero E6 cells by SARS-CoV-2-, SARS-CoV-, and MERS-CoV-S-pseudotyped virions

Previous studies had shown that CMC-Cu nanoparticles exhibited antimicrobial properties as a result of inhibition of bacterial and fungal growth ^26, 28, 29^. To test whether treatment with CMC-Cu affected viability of Vero E6 cells, the cells were exposed (30 s) to a concentrated CMC-Cu solution (100%) as well as a series of diluted CMC-Cu concentrations over a range from 0.1 to 75%. Cell viability was determined by MTT assays after a 72-h period of incubation. Results showed that the CMC-Cu concentration found to foster 50% inhibition (IC_50_) of viability was 9.93%. In contrast, CMC-Cu used at a concentration lower than 4% did not affect Vero E6 cell viability (Fig. S1, B). To evaluate the virucidal effect of CMC-Cu on SARS-CoV-2-, SARS-CoV-, and MERS-CoV-S-pseudovirions, the S-pseudotyped virions were exposed to concentrations of CMC-Cu ranging from 0.0005 to 4% over a period of time of 30 s. The virus-like particles were assayed for their ability to infect Vero E6 cells after 72 h. Results showed that the concentration of CMC-Cu needed for half maximal inhibition (IC_50_) of the infectivity of SARS-CoV-2-, SARS-CoV- and MERS-CoV-S-pseudovirions was 0.09%, 0.11%, and 0.45%, respectively (Fig. 4, A – C). Interestingly, these results revealed that SARS-CoV-2-, SARS-CoV- and MERS-CoV-S-pseudoviral particles were 110-, 90-, and 22-fold more sensitive to CMC-Cu nanoparticles than Vero E6 cells.

**Fig. 4.**
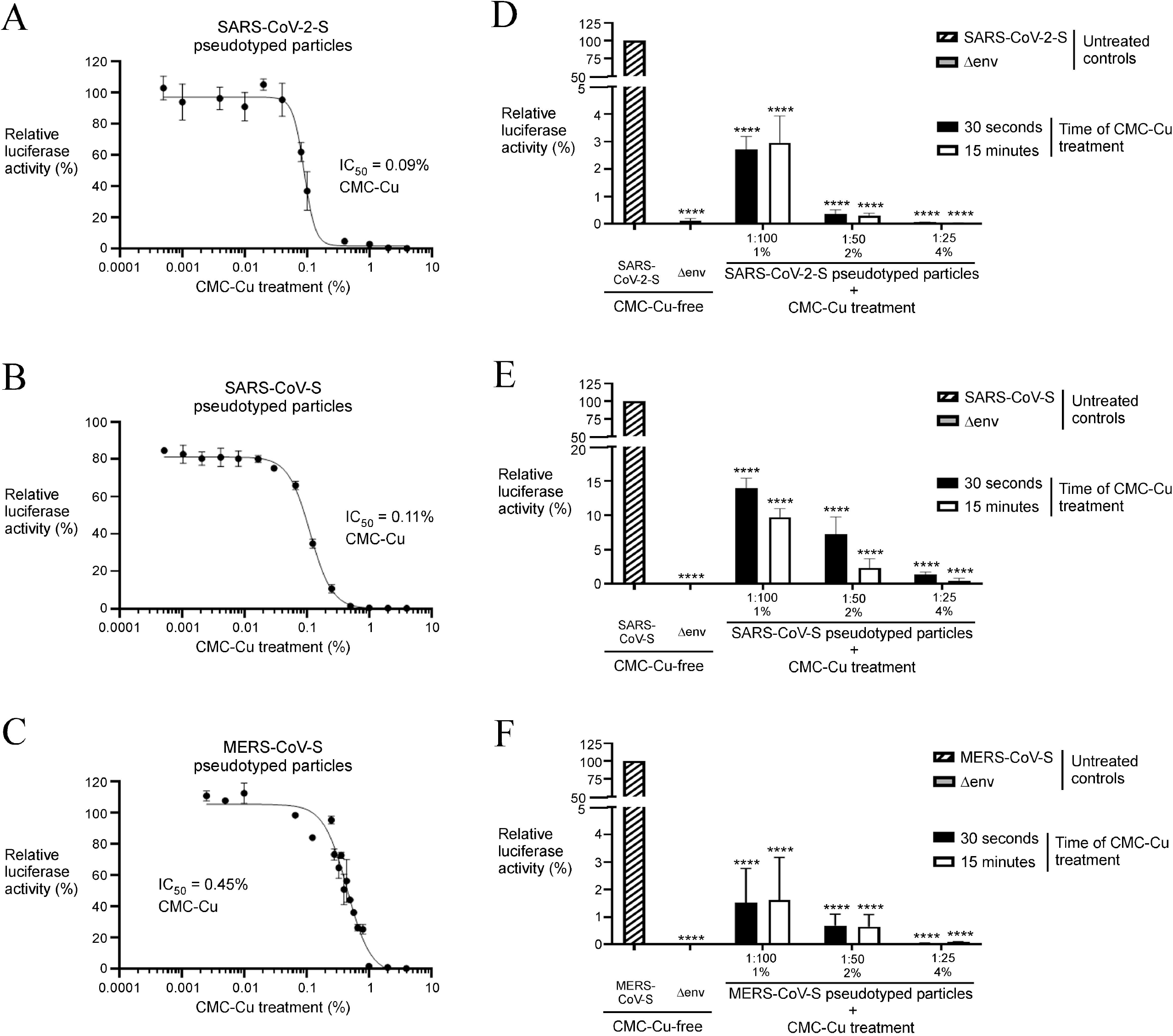
CMC-Cu inhibits infectivity of SARS-CoV-2-, SARS-CoV-, and MERS-CoV-S-pseudotyped virions. *A – C*, The S-pseudovirions were treated with CMC-Cu concentrations from 0.0005 to 4% for 30 s. The S-pseudovirions were then used to infect Vero E6 cells and quantification of host cell susceptibility was performed by luciferase assays 72 h after infection. *D – F*, SARS- CoV-2-, SARS-CoV-, and MERS-CoV-S-pseudotyped virions were exposed to CMC-Cu at diluted concentrations of 1:100, 1:50, and 1:25 for 30 s or 15 min prior to Vero E6 cell infection. As controls, Vero E6 cells were infected with untreated S-pseudovirions (CMC-Cu-free, positive control) or pseudovirions lacking a cell-surface viral envelope glycoprotein (Δenv, negative control). Luciferase assays were carried out 72 h post infection. Results of luciferase assays of cells with untreated S-pseudovirions was set up to 100%. Graphs represent quantification of the results of at least five independent infection experiments performed in biological triplicate. Values are represented as averages ± S.D. The asterisks correspond to *p* < 0.0001 (****) (one-way ANOVA with Dunnett’s multiple comparisons test against the CMC-Cu-free positive control).

Concentrations of CMC-Cu that were effective in inactivating S protein-bearing pseudoviral particles but that were non-toxic to Vero E6 cells were selected for a next series of experiments. S-pseudovirions were pretreated with the CMC-Cu solution at diluted concentrations of 1:100 (1%), 1:50 (2%), and 1:25 (4%) that corresponded to CMC containing reduced Cu at concentrations of 200, 400, and 800 µM, respectively. Pretreatment of S-pseudovirions were performed over a period of time of 30 s or 15 min prior to assays of Vero E6 cells infection. Results showed that cells that had been exposed to CMC-Cu-treated SARS-CoV-2-S pseudoviral particles (CMC-Cu at a concentration of 1:100) displayed 97.1% to 97.3% less luciferase activity compared to the untreated pseudoviral particles (Fig. 4D). Furthermore, we observed a decrease of 99.6% to 99.9% less luciferase activity when Vero E6 cells were infected with pseudoviral particles expressing SARS-CoV-2 S protein that had been pretreated with CMC-Cu at a dilution of 1:50 and 1:25, respectively (Fig. 4D).

In the case of SARS-CoV-S pseudovirions, a pretreatment with CMC-Cu (1:100) resulted in a strong reduction in their ability to infect Vero E6 cells as shown by a 86.1% to 90.3% reduced luciferase activity (Fig. 4E). In a next series of experiments, SARS-CoV-S pseudovirions were treated with CMC-Cu at concentration of 1:50 and 1:25 and then incubated (30 s and 15 min) with target Vero E6 cells. Results showed efficient inhibitory effect of cell infection as shown by a decrease of 92.8% to 99.5% of luciferase activity compared to the absence of CMC-Cu treatment (Fig. 4E).

Pseudovirions expressing the MERS-CoV-S protein were treated with CMC-Cu at the indicated diluted concentrations (1:100, 1:50 and 1:25) for 30 s and 15 min and assayed for their ability to infect Vero E6 cells. Luciferase activity was then measured 72 h post treatment. Results showed a reduction of infectivity of 98.4%, 99.4%, and 99.9% in comparison with the absence of CMC-Cu treatment (Fig. 4F). To analyze data, the luciferase activity of infected cells with MERS-CoV- S (or SARS-CoV-S or SARS-CoV-2-S) pseudovirions without CMC-Cu treatment was set up to 100% and luciferase activities of cells infected with CMC-Cu-treated S-pseudovirions were measured and compared to untreated S-pseudovirions. Taken together, these results showed that CMC-Cu possesses antiviral properties that are effective to inhibit the infectivity of SARS-CoV- 2-, SARS-CoV-, and MERS-CoV-S-pseudovirions.

### Treatment with Saber^TM^ in combination with CMC-Cu is the most effective way to restrict infectivity of SARS-CoV-2-, SARS-CoV-, and MERS-CoV-S pseudovirions

We next sought to determine whether a combination of Saber^TM^ (0.4%) and CMC-Cu (at diluted concentration of 1:100; containing 200 µM Cu) would result in a stronger reduction of the luciferase signal. In the case of SARS-CoV-2-S pseudovirions, treatment (30 s) with Saber^TM^ alone resulted in a 97.6% reduction of the luciferase signal compared to control conditions (absence of treatment) (Fig. 5A). When CMC-Cu alone was used to treat (30 s) SARS-CoV-2-S pseudovirions prior Vero E6 infection, the luciferase signal was reduced to 92.1% (Fig. 5A). Treatment of SARS-CoV-2-S pseudovirions under similar conditions with a combination of Saber^TM^ (0.4%) and CMC-Cu (1:100) resulted in near background values of luciferase activity (99.8% reduction) (Fig. 5A).

**Fig. 5.**
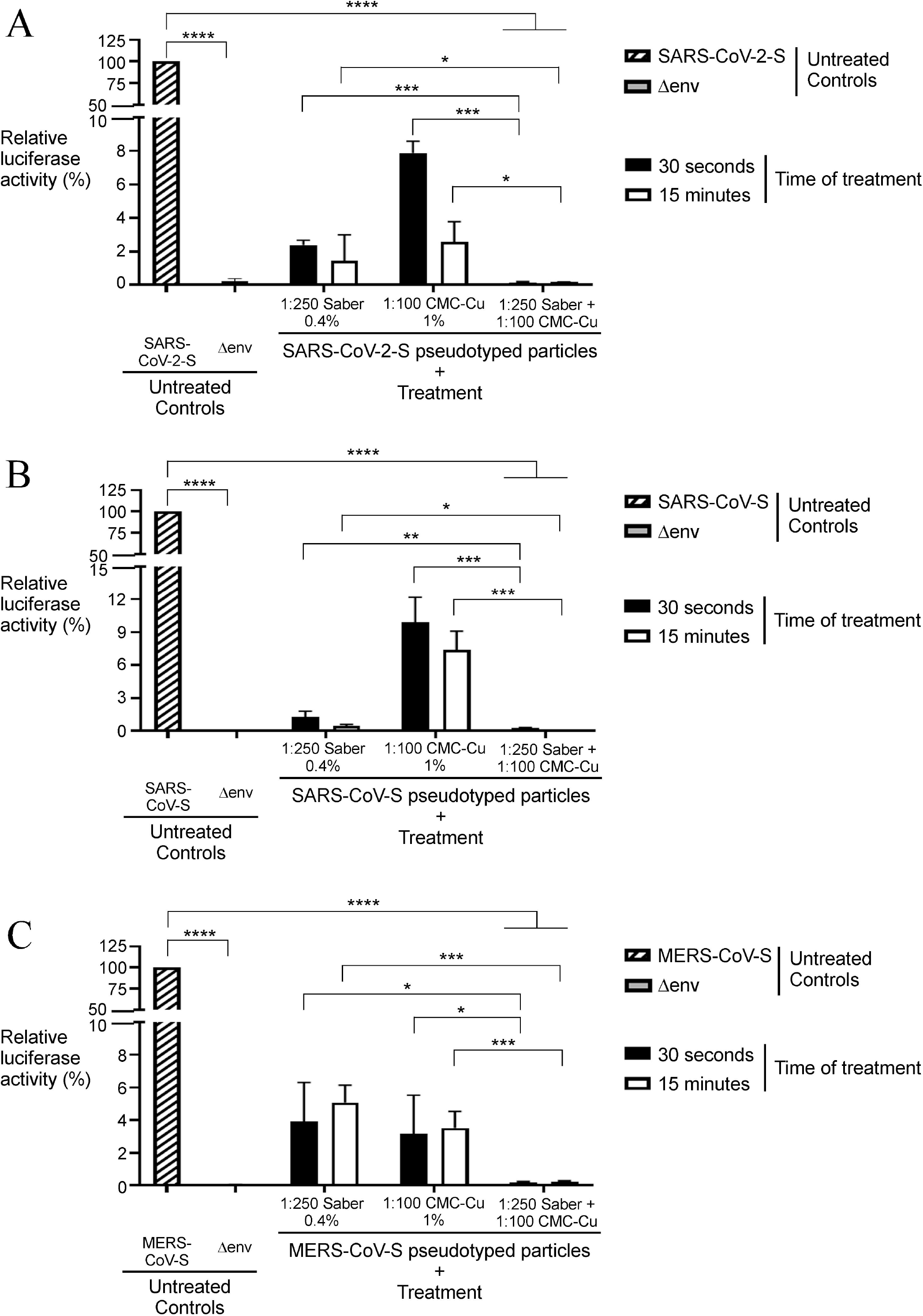
Maximal inhibition of MLV-based pseudovirions expressing the S protein requires the combination of Saber^TM^ and CMC-Cu. *A – C*, SARS-CoV-2-, SARS-CoV-, and MERS-CoV-S- pseudotyped virions were treated with Saber^TM^ (1:250), CMC-Cu (1:100) or Saber^TM^ in combination with CMC-Cu for 30 s or 15 min prior to Vero E6 cell infection. As controls, Vero E6 cells were infected with untreated S-pseudoviral particles (Saber^TM^- and CMC-Cu-free, positive control) or pseudovirions lacking a cell-surface viral envelope glycoprotein (Δenv, negative control). Luciferase activity was measured 72 h post infection. Results of luciferase activity of infected cells with untreated S-pseudovirions was set up to 100%. Graphs represent quantification of a minimum of five independent experiments that were performed in biological triplicate. Values are represented as averages ± S.D. The asterisks correspond to *p* < 0.05 (*), *p* < 0.01 (**), *p* < 0.001 (***), and *p* < 0.0001 (****) (one-way ANOVA with Dunnett’s multiple comparisons test against the untreated S-pseudovirions or under the conditions in which Saber^TM^ and CMC-Cu were used separately to treat S-pseudovirions). ns stands for not significant.

Treatment of SARS-CoV-S-pseudovirions with a combination of Saber^TM^ and CMC-Cu resulted in reduction of 99.7% to 99.9% of luciferase activity in target Vero E6 cells compared to control experiments (untreated pseudovirions) (Fig. 5B). When Saber^TM^ and CMC-Cu were used separately to treat pseudovirions expressing the SARS-CoV-S protein, the luciferase signal observed in cell lysates was reduced by 98.7% and 90.1%, respectively (Fig. 5B). In other sets of experiments, Saber^TM^ was used in combination with CMC-Cu to treat MERS-CoV-S-bearing pseudovirions. Results showed a dramatically reduced luciferase activity that was 99.8% less compared to the untreated pseudoviral particles (Fig. 5C). Treatment of MERS-CoV-S pseudovirions with Saber^TM^ or CMC-Cu resulted in 96.1% and 96.5% inhibition of luciferase activity, respectively, compared to the absence of treatment (Fig. 5C). Taken together, these results revealed that a treatment of SARS-CoV-2-, SARS-CoV-, and MERS-CoV-S pseudovirions with a combination of the Saber^TM^ and CMC-Cu is the most efficient way to inactivate their ability to infect.

### The Cu chelator TTM and the ROS quencher Tiron protect S-pseudoviral particles against the inhibitory effect of CMC-Cu treatment

Results described above showed very similar results whether either one or the other MLV-based coronavirus S-pseudotyped virus was tested. Therefore, we only used the SARS-CoV-2-S-pseudotyped virus for the next series of experiments. The redox active nature of Cu makes it highly susceptible to engage in chemical reactions that generate hydroxyl radical, a free radical species that damages membrane lipids, proteins and nucleic acids. These observations led us to test whether the Cu chelator tetrathiomolybdate (TTM) would protect S-pseudovirions against the CMC-Cu nanotoxicity. As a control, results showed that incubation of S-pseudoviral particles with TTM (4 mM) alone did not affect their ability to infect Vero E6 cells. Indeed, lysates from these cells yielded high levels of luciferase activity that were set to 100% compared with low background levels of luciferase activity (0.2%) in cell extracts from cells infected with Δenv pseudoviral particles (negative control) (Fig. 6A). When SARS-CoV-2-S pseudovirions were treated with CMC-Cu (1:100) over a period of time of 30 s, there was a strong inhibitory effect on their infectivity. Luciferase assays showed a 97.6% reduction of activity compared to the absence of treatment (Fig. 6A). In contrast, when S-pseudovirions were treated with CMC-Cu (1:100) and TTM (4 mM), levels of luciferase activity were restored to 95.6% compared to levels of S-pseudoviral particles not exposed to CMC-Cu (Fig. 6A).

**Fig. 6.**
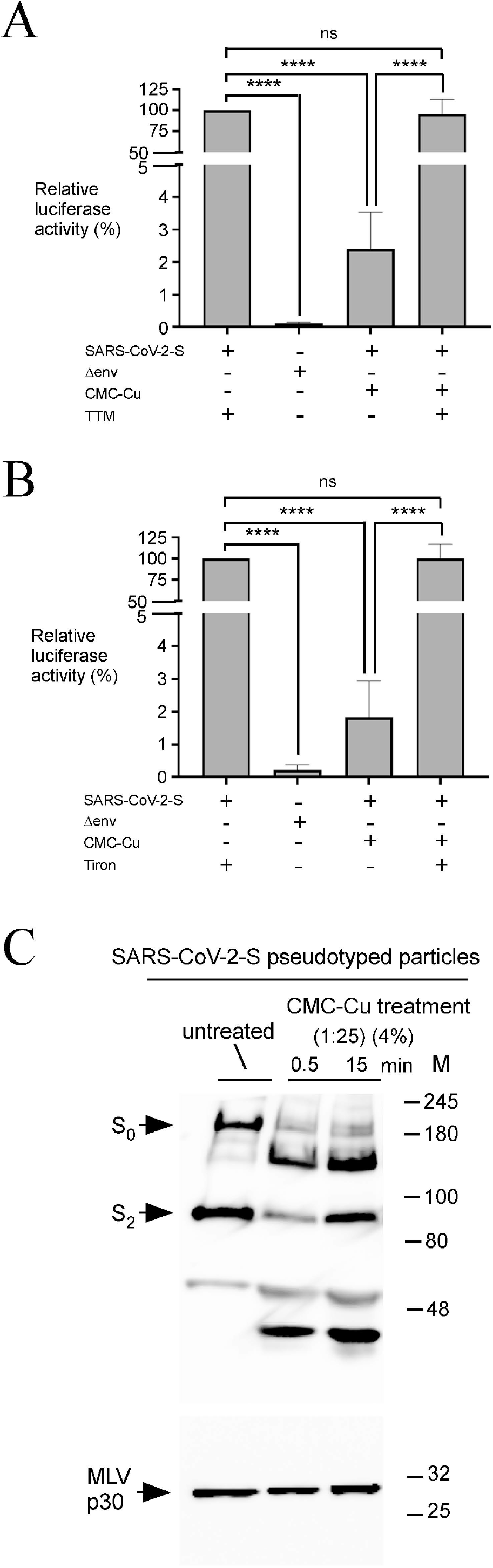
TTM and Tiron protect SARS-CoV-2 S-pseudovirions against the inhibitory effect of CMC-Cu. *A*, SARS-CoV-2 S-pseudotyped virions were incubated in the presence of TTM (4 mM), CMC-Cu (1:100), or CMC-Cu in combination with TTM over a period of 30 s prior to Vero E6 cell infection. As a negative control, Vero E6 cells were infected with pseudovirions lacking the cell-surface viral envelope S protein (Δenv). *B*, Aliquots of the S-pseudovirions were treated with Tiron (100 mM), CMC-Cu (1:100), or CMC-Cu in combination with Tiron for 30 s prior to Vero E6 cell infection assays. Luciferase activity was measured 72 h post infection. The luciferase signal of cells infected by S-pseudovirions that had been treated with TTM or Tiron alone was set up to 100%. Graphs represent quantification of the results of a minimum of five independent experiments performed in biological triplicate. Values are represented as averages ± S.D. The asterisks correspond to *p* < 0.0001 (****) (one-way ANOVA with Dunnett’s multiple comparisons test against the untreated SARS-CoV-2-S-pseudotyped particles, unless otherwise stated), whereas ns stands for not significant. *C*, Aliquots of untreated or CMC-Cu-treated S-pseudotyped particles were analyzed by immunoblot assays using anti-SARS-CoV-2 Spike and anti-MLVp30 antibodies. The positions of the molecular weight standards (in kDa) are indicated on the right side.

To further investigate the mechanism of the action of CMC-Cu, we investigated the protective effect of Tiron, a hydroxyl radical quencher, to interfere with the inhibitory action of CMC-Cu nanoparticles. As described above, a treatment with CMC-Cu (1:100) within 30 s strongly inhibited the ability of S-pseudovirions to infect Vero E6 cells, resulting in luciferase activity that was 98.2% less than activity of untreated S-pseudovirions (CMC-Cu free) (Fig. 6B). When Tiron and CMC-Cu were used together to treat pseudovirions expressing the SARS-CoV-2-S protein, the luciferase signal observed in Vero E6 cells was fully restored to 100% of that of reference S- pseudovirions (Fig. 6B). Taken together, these results suggested that the redox active nature of Cu was required to inactivate S-pseudovirions.

To further investigate the relationship between CMC-Cu and S-pseudoviral particles, Western blot assays were performed to examine the S protein levels of purified pseudovirions following their treatment with CMC-Cu after 30 s and 15 min. Results showed that the levels of S_0_ form of Spike were rapidly reduced when pseudovirions had been incubated with 4% CMC-Cu (dilution 1:25) (Fig. 6C). Moreover, additional low molecular weight bands were present and increased in intensity as a function of time of treatment (Fig. 6C). In the case of the S_2_ form, protein fragment levels appeared relatively stable in the presence of CMC-Cu. The capsid MLV p30 protein levels were probed as loading control. It is tempting to suggest that the rapid loss of S_0_ form integrity of the S protein may contribute to the dramatic reduction of infectivity of viral pseudotyped particles exposed to CMC-Cu.

### Combined treatment with Saber^TM^ and CMC-Cu inhibits SARS-CoV-2 S-pseudotyped variants to the same extent as pseudoviral particles expressing wild-type S protein

Since the onset of the COVID-19 pandemic, several studies have shown a constant emergence of genomic sequence variations of SARS-CoV-2 isolates.^50–53^ Several of these genomic mutations have been identified in the gene encoding the S protein.^54^ One mutation results from substitution of adenine by guanine at position 23,403 in the genome of the Wuhan reference strain. This substitution produces a new variant containing an aspartic acid to glycine substitution at amino acid residue 614, called D614G.^55^ Another example of mutation consists in adenine to thymine substitution at position 23,063. In this case, the substitution generates a mutant containing an asparagine to tyrosine at amino acid residue 501, denoted N501Y.^56^ To further evaluate the efficiency of Saber^TM^ + CMC- Cu combination to block infectivity of SARS-CoV-2-S-pseudotyped variants, we produced the following mutant versions of S protein-bearing pseudoviral particles: 501Y, 614G, and 501Y/614G.

We first used equal amounts of pseudoviral particles expressing different versions of S protein to infect Vero E6 cells. Results showed that Vero E6 cells infected with pseudovirions expressing the mutated S 501Y/614G protein exhibited the highest levels of luciferase activity (1.5 x 10^5^ RLU) compared to levels of cells that had been infected with pseudoviral particles expressing the wild-type S protein (8.9 x 10^4^ RLU) (Fig. 7A). This observation suggested that Vero E6 cells were highly permissive to infection by pseudovirions expressing the S 501Y/614G protein. When S 501Y and S 614G mutant proteins were individually expressed on the surface of pseudovirions and used to infect Vero E6 cells, luciferase activity increased to 1.2 x 10^5^ and 1.4 x 10^5^ RLU, respectively, compared to levels of pseudoviral particles that expressed wild-type S protein (8.9 x 10^4^ RLU) (Fig.7A). However, their average values of luciferase activity in these instances were lower than in the case of cells infected with pseudoviral particles expressing the S 501Y/614G protein (Fig. 7A).

**Fig. 7.**
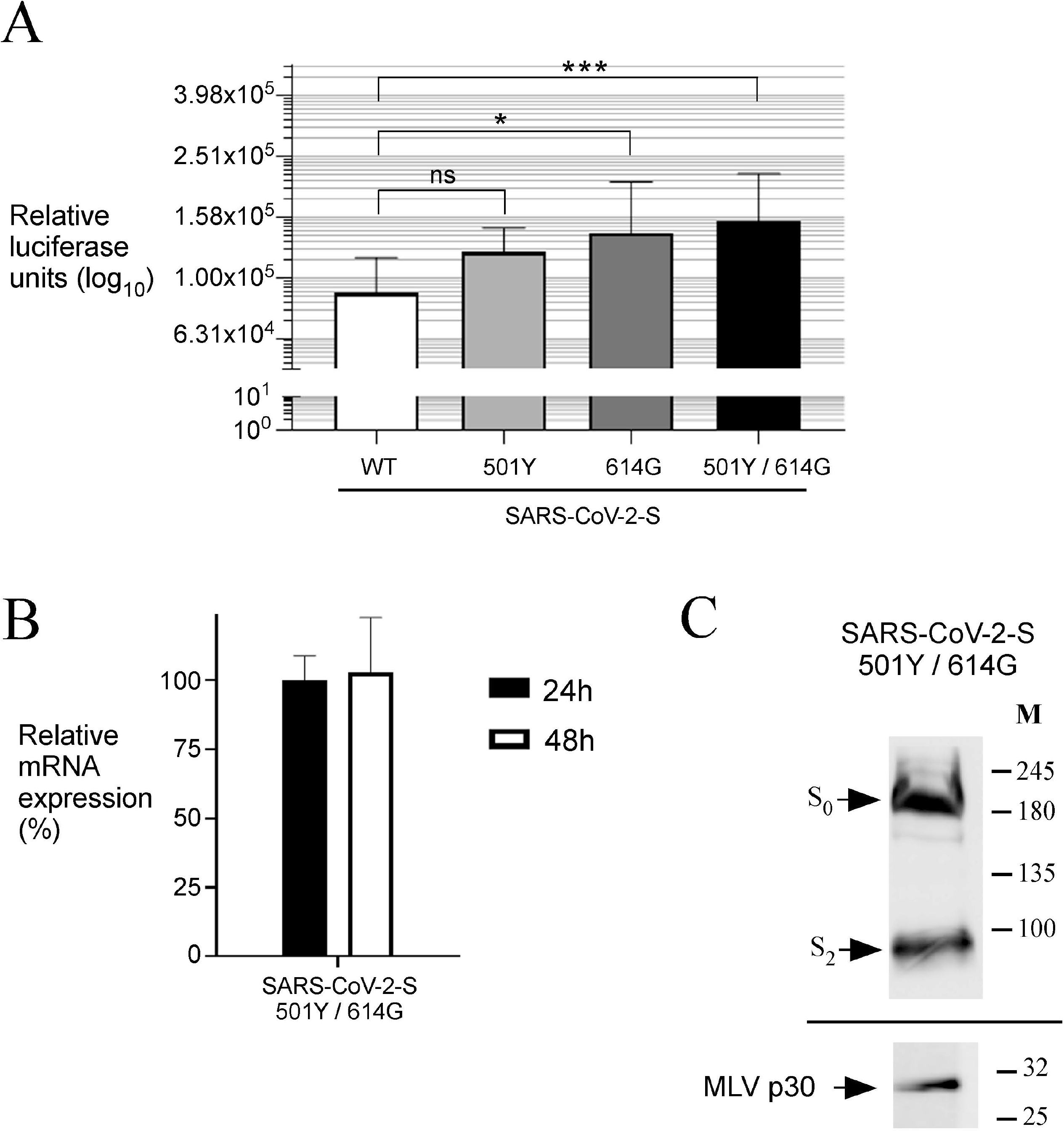
Infection of Vero E6 cells by wild-type and mutant SARS-CoV-2 S protein-pseudotyped MLV viral particles. *A*, Vero E6 cells were infected by the indicated wild-type or mutant S protein-expressing MLV-based pseudotyped particles. Data presented are average relative luciferase units (log_10_) of five independent experiments 72 h post infection (performed in biological triplicate). Pseudoviral particle codes are as follows: WT, wild-type; 501Y, the N501Y mutant of SARS-CoV-2 Spike; 614G, the D614G mutant of SARS-CoV-2 Spike; 501Y/614Y, the N501Y and D614G double mutant of SARS-CoV-2 Spike. Values are represented as averages ± S.D. The asterisks correspond to *p* < 0.05 (*) and *p* < 0.001 (***) (one-way ANOVA with Dunnett’s multiple comparisons test against wild-type SARS-CoV-2-S-pseudotyped particles). ns stands for not significant. *B*, Total RNA was isolated from HEK293T cells that had been cotransfected with plasmids encoding the luciferase gene, MLV Gag-Pol loci, and the SARS-CoV- 2-S gene harboring mutations at positions 23,063 and 23,403, generating amino acid substitutions N510Y and D614G in Spike protein. Following RNA isolation at the indicated time points, SARS-CoV-2-S 501Y/614G mRNA levels were analyzed by RT-qPCR assays. A first primer set was used to probe an unique region of SARS-CoV-2-*S* 501Y/614G mRNA, whereas a second primer set was used to detect human B2M transcript as an internal control for normalization during quantification of the RT-qPCR products. *C*, Western blot analysis of SARS-CoV-2-S 501Y/614G and MLV p30 (control) levels after 48 h transfection. S_0_, uncleaved form of S protein; S_2_, S2 segment of S protein. The positions of the molecular weight standards (in kDa) are indicated on the right side.

During pseudotyped particle production in HEK-293T cells, we validated that S mutant proteins of SARS-CoV-2 were expressed at both transcriptional and posttranscriptional levels. We focused our attention on the S 501Y/614G mutant since pseudoviral particles expressing this variant form of S protein gave the highest luciferase signal. As shown in Fig. 7B, S 501Y/614G RNA transcripts were strongly detected in total RNA preparations from HEK-293T after 24 and 48 h of transfection. When pseudotyped particles were collected at 48 h time point, results showed that the S 501Y/614G mutant protein was consistently detected from pseudovirions (Fig. 7C). The two forms of S 501Y/614G that were detected included the uncleaved (S_0_) form and the S_2_ segment (S_2_) of the protein as reported previously (Fig. 7C).^48, 49^

To assess the efficiency of Saber^TM^ and CMC-Cu against S 501Y/614G pseudoviral particles, we incubated aliquots of these pseudovirions with Saber^TM^ and CMC-Cu concentrations ranging from 0.0004 to 1% and 0.0005 to 4%, respectively. After 30 s, Saber^TM^- and CMC-Cu-treated S 501Y/614G pseudotyped particles were used to test Vero E6 cells infection by luciferase assays at 72 h (Fig. 8, A – B). Results showed that half maximal inhibitory concentration (IC_50_) of Saber^TM^ and CMC-Cu to block the infectivity of S 501Y/614G pseudotyped particles was 0.02% and 0.06%, respectively (Fig. 8, A – B). These results indicated that viral pseudotyped particles were 74- and 170-fold more sensitive to Saber^TM^ and CMC-Cu, respectively, than Vero E6 cells.

**Fig. 8.**
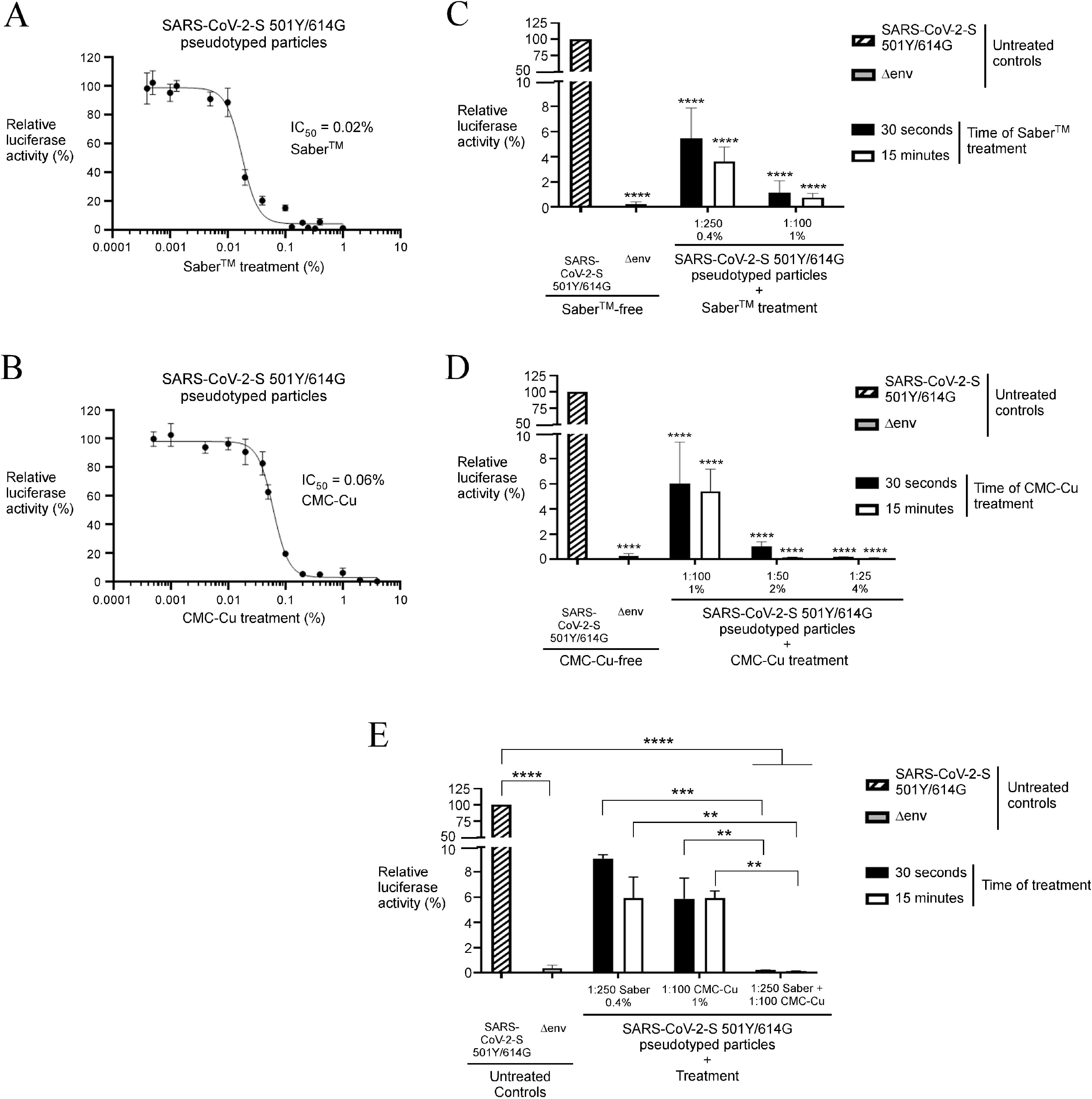
Pseudovirions containing wild-type or mutant S proteins of SARS-CoV-2 are more effectively inactivated by the combined use of Saber^TM^ with CMC-Cu. *A – B,* aliquots of S 501Y/614G pseudotyped particles were treated with the indicated concentrations of Saber^TM^ (*panel A*) or CMC-Cu (*panel B*) for 30 s. The pseudovirions were then used to infect Vero E6 cells and quantification of cell entry was performed by luciferase assays*. C – E*, pseudoviral particles expressing the S 501Y/614G mutant protein were treated with Saber^TM^ (0.4%, 1:250; or 1%, 1:100) (*panel C*). Separate aliquots of these pseudoviral particles were also treated with different concentrations of CMC-Cu at diluted concentrations of 1:100, 1:50, and 1:25 (*panel D*). The last treatment consisted of incubating pseudoviral particles in the presence of Saber^TM^ (1:250) in combination with CMC-Cu (1:100) (*panel E*). All these treatments were performed over a period of 30 s or 15 min prior to Vero E6 cell infection. As controls, Vero E6 cells were infected with untreated pseudoviral particles (Saber^TM^- and CMC-Cu-free, positive control) or pseudovirions lacking a cell-surface viral envelope glycoprotein (Δenv, negative control). Luciferase activity was measured 72 h post infection. Values of luciferase assays of infected cells with untreated S 501Y/614G pseudovirions were set up to 100%. Graphs represent quantification of the results of a minimum of five independent experiments performed in biological triplicate. Values are represented as averages ± S.D. The asterisks correspond to *p* < 0.01 (**), *p* < 0.001 (***), and *p* < 0.0001 (****) (one-way ANOVA with Dunnett’s multiple comparisons test against the untreated SARS-CoV-2-S-501Y/614G pseudotyped particles or in the case of *panel E*, under the conditions in which Saber^TM^ and CMC-Cu were used separately to treat pseudotyped particles).

Based on these results, we selected Saber^TM^ and CMC-Cu concentrations that were effective in inactivating S 501Y/614G pseudovirions but that non-toxic to Vero E6 cells. Pseudoviral particles expressing S 501Y/614G were treated with Saber^TM^ at concentrations of 1:250 (0.4%) and 1:100 (1%) for 30 s or 15 min. Results showed that Vero E6 cells exposed to Saber^TM^-treated S 501Y/614G pseudovirions exhibited 94.5% to 96.4% and 98.8% to 99.3% less luciferase activity (depending on the concentration of Saber^TM^ used) compared to the absence of Saber^TM^ (Fig. 8C). When S 501Y/614G pseudotyped particles were pretreated with CMC-Cu at concentrations of 1:100, 1:50, and 1:25 over a period of time of 30 s or 15 min, there was also a strong inhibition in their capacity to infect Vero E6 cells, exhibiting 94.0% to 99.9% less luciferase activity as a function on the increasing concentration of CMC-Cu compared to absence of CMC-Cu treatment (Fig. 8D). These results led us to select the lowest concentrations of Saber^TM^ (1:250) and CMC- Cu (1:100) to test whether their combinations would result in a greater inhibitory effect to restrict infectivity of pseudoviral particles expressing S 501Y/614G. Treatment of pseudovirions for 30 s and 15 min with Saber^TM^ alone (1:250) resulted in 90.9% to 94.1% less luciferase activity, whereas treatment with CMC-Cu alone (1:100) inhibited the luciferase signal by 94.1% compared to untreated pseudovirions (Fig. 8E). Importantly, treatment of S 501Y/614G pseudotyped particles with Saber^TM^ (1:250) in combination with CMC-Cu (1:100) resulted in the most efficient way to block their entry into Vero E6 cells in luciferase assays. Data showed a dramatic reduction to background values with the lowest signal intensity (99.9% reduction) (Fig. 8E). In contrast, control experiments revealed maximal luciferase activity in cell extracts prepared from Vero E6 cells infected with untreated S 501Y/614G pseudotyped particles. Taken together, these results revealed that Saber^TM^ in combination with CMC-Cu remain highly effective to inhibit SARS-CoV- 2 S-pseudotyped particle variants harboring 501Y, 614G, and 501Y/614G mutations.

## Discussion

Although SARS-CoV-2 infections mostly occur as a result of person-to-person contact through exposure to respiratory droplets, viral transmission can also occur from high-touch contaminated surfaces to hands, and from hand-to-face contacts as observed for other pathogens.^57–59^ Previous studies have shown that disinfection of high-touch non-porous surfaces can reduce the risk of fomite transmission for SARS-CoV-2.^10, 60^ Copper-based surfaces and Cu alloys containing at least 70% Cu are effective at inactivating bacteria and viruses.^21–23^ The antimicrobial properties of Cu-based surfaces rely on the release of deleterious redox active Cu ions from the surface. Progressive Cu release triggers severe damages to the lipid bilayer envelope of bacteria and viruses. The loss of membrane integrity allows increased influx of Cu ions into the cell or viral particle that causes detrimental oxidative damages to lipids, proteins, and nucleic acids. Although the successive steps leading to microbial cell death may vary upon the type of microorganism, microbe-Cu contacts are required for Cu-based surface mediated killing.^25^

Based on this knowledge, we used a carboxymethyl cellulose (CMC) nanostructure as a physical template to form and stabilize Cu-containing nanoparticles that can be use as an antimicrobial nanocomposite against enveloped MLV-based pseudovirions expressing the S protein of SARS-CoV-2, SARS-CoV, and MERS-CoV. CMC-Cu nanostructure possesses Cu-containing nanospheres on its surface of sizes ranging from 20 nm to 2000 nm. This structure corresponds to a surface sufficient to allow physical interaction with a MLV-based pseudovirus that has a diameter of 80 to 150 nm.^26, 27, 61–63^ In this way, the antimicrobial potency of Cu could be increased by the delivery of Cu in the form of CMC-Cu, therefore creating a “copperized nanosurface” that favors direct pseudovirus-Cu contacts, rendering Cu more effective at inactivating pseudovirions. In support of this hypothesis, a previous work has shown that yeast cells exposed to CMC-Cu containing the equivalent of 157 µM Cu exhibit the same level of growth inhibition than the same cells exposed to soluble Cu in which case 400 µM of cupric sulfate is present, revealing that Cu exhibits a greater toxicity under the form of CMC-Cu than soluble Cu.^28^ In addition, a report has shown that carboxymethyl cellulose (CMC) is the best support for catalyst Cu as it incorporates the highest amount of Cu per gram of nanomaterial due to its higher number of sodium-carboxyl groups.^64^ Furthermore, analyses have shown that the oxidative state of Cu in the final nanomaterial product is Cu^1+^, making it able to engage in chemical reactions that generate hydroxyl radical, a free radical species that is cytotoxic.

In the case of pseudoviral particles used in our study, they consisted of a surrogate viral core with a heterologous coronavirus S protein at their surface. Previous studies have shown that these pseudoviral particles behave like their native coronavirus counterparts for entry steps into susceptible cells.^36, 38^ Thus, use of these S-expressing pseudoviral particles represents an excellent model for investigating viral entry into host cells. A 30 s or 15 min of treatment with CMC-Cu was performed to test whether cellulosic Cu nanoparticles have an impact on pseudovirion entry mediated by the S protein. A treatment of 30 s was used in accordance with ASTM International standard (E-1052) ^65^ with respect to hand hygiene practices against enveloped viruses such as SARS-CoV-2. The second treatment of 15 min was performed to verify whether the virucidal effect gave results similar to a 30-s treatment. Results showed that SARS-CoV-2-, SARS-CoV-, and MERS-CoV-S-pseudotyped particles were extensively inactivated by CMC-Cu within 30 s of exposure. A treatment with CMC-Cu at a diluted concentration of 1:100 resulted in 86% to 98% decrease of luciferase activity, whereas S-pseudovirions infectivity was virtually abolished (≥99%) when they have been treated with the CMC-Cu solution at a concentration of 1:25. These results correspond to a clear loss in viral infectivity, with a 1.6 log_10_ reduction at a dilution of 1:100, a 2.4 log_10_ reduction at a dilution of 1:50, and a 3.2 log_10_ reduction at a dilution of 1:25 in the case of SARS-CoV-2-S pseudovirions (Fig. S2). Furthermore, a 30-s treatment for inactivation of pseudoviral particles was as effective as a 15-min treatment. For comparison purpose, it has been previously determined that the human coronavirus 229E was completely inactivated after 20 min of contact with a brass surface containing 90% Cu, exhibiting a 3.0 log_10_ reduction in virus concentration.^23^ However, analysis of early contact time points (5- and 10-min exposure) revealed a lag in inactivation of about 10 min followed by a rapid loss of infectivity.^23^ Based on our results, the advantage of using CMC-Cu resides in the fact that it would offer a faster virucidal effect of Cu on S-pseudoviral particles, triggering their inactivation after only 30 s.

We used the H_2_O_2_-based disinfectant Saber^TM^ at concentrations of 1:250 and 1:100 to compare the efficacy of CMC-Cu for inactivation of S-expressing pseudovirions. In addition to containing H_2_O_2_, Saber^TM^ includes the anionic surfactant sodium dodecylbenzenesulfonate to improve its virucidal efficiency. After the exposure of pseudovirions (30 s or 15 min) to Saber^TM^, the mixture was filtered twice through a Sephadex LH-20 column in accordance with ASTM International standard (E-1482) ^66^ to minimize any potential cytotoxic effect on Vero E6 cells. The filtrate (containing S-pseudoviral particles) was assayed on Vero E6 cells. Results showed that SARS-CoV-2-, SARS-CoV-, and MERS-CoV-S-pseudotyped particles were sensitive to Saber^TM^ for a contact time of 30 s. This time of exposure reflects the habits of customers when they disinfect their hands before entering a commercial outlet. S-pseudoviral particles infectivity was decreased by 98.7% to 99.4% when they have been treated with 0.4% Saber^TM^. For one particular example, when SARS-CoV-2-S pseudovirions were exposed to 0.4% and 1% of Saber^TM^ for 30 s, ≥2.2- and 2.5-log_10_ inactivation was observed (Fig. S3). These drops in reporter activity revealed that the most of the infectivity of S-pseudoviral particles was eliminated by 0.4% and 1% of Saber^TM^ exposure of 30 s. These results were reminiscent of the effective inactivation of human coronaviruses (HCoV) by disinfection procedures with 70% EtOH, 70% isopropanol, 0.5% H_2_O_2_, or 0.1% sodium hypochlorite within a 30 s or 60 s exposure.^30, 67^

Based on a previous study that reported enhanced virucidal effect of Cu by the addition of hydrogen peroxide ^68^, we have investigated whether a treatment with CMC-Cu in combination with peroxide-based Saber^TM^ potentiated the level of inactivation of S-pseudoviral particles. Results showed that S-pseudovirions were more sensitive to the Saber^TM^-CMC-Cu mixture than to Saber^TM^ or CMC-Cu alone. In the case of SARS-CoV-2 S-pseudotyped particles, a treatment with the Saber^TM^-CMC-Cu mixture resulted in a 2.8-log_10_ decrease which represented 100% reduction compared to background levels of infectivity observed with Δenv pseudoviral particles in Vero E6 cells assays (Fig. S4).

A previous study has reported that 10- or 30-min exposure to a Cu brass surface resulted in morphological alterations to the human coronavirus 229E, including envelope damages and loss of the surface S protein.^23^ Because the Cu^1+^ chelator bathocuproine disulfonate (BCS) protected HuCoV-229E on Cu brass surface, it has been proposed that release of Cu^1+^ from the surface material was required for the virucidal effect and supporting the generation of destructive hydroxyl radicals via the Fenton reaction. The results reported here suggest that a similar Cu^1+^-dependent ROS-catalyzed killing occurred in the presence of the nanoparticles since the Cu^1+^ chelator TTM and hydroxyl radical quencher Tiron protected S protein-bearing pseudovirions against CMC-Cu. Nonetheless of the incompletely understood biocidal mechanisms of CMC-Cu, its use without or optimally in combination with a peroxide-based disinfectant could help to reduce infection spread from touching surfaces contaminated with enveloped coronaviruses.

## Data availability

All data are included in the present manuscript. Strains and plasmids used for this study are available upon request. The authors state that all results obtained for confirming the conclusions presented in the article are represented fully within the article.

## Author contributions

A.B., S.B., M.B., P.M., P.C., and S.L. conceptualized the experimental work. A.B. designed and modified DNA constructs. A.B., R.N., J.P., S.B. performed experiments. A.B., R.N., J.P., S.B., M.B., and S.L. acquired and analyzed the data. A.B. and S.L. wrote the manuscript. The authors reviewed the results and approved the final version of the manuscript.

## Conflict of interest – Disclaimer statement

PM and PC are employed with Sani-Marc Group Inc. and have no financial interest in the subject matter or information presented and discussed in this manuscript. The authors declare that they have no known personal relationships that could have appeared to influence the work reported in this study. The other authors declare that they have no conflict of interest with the content of this article.

## Acknowledgments

We are grateful to Dr. Gilles Dupuis for critical reading of the manuscript and for his valuable comments. We are thankful to Sébastien Dion, Antoine Désilets, Wendy O. Wingate, and Drs. Jean Kaoru Millet, Gary R. Whittaker, and Richard Leduc for their generous gift of cell line and plasmids. This study was supported by the *Ministère de l’Économie et de l’Innovation* (MÉI, grant 2020-2021-COVID-19-PSOv2a-51192) and, in part, by the Natural Sciences and Engineering Research Council of Canada (NSERC, grant #ALLRP 554545-20) to S. Labbé.

## Footnotes

^1^Abbreviations used are: ASTM, American Society for Testing and Materials; B2M, human β_2_-microglobulin; bp, base pair(s); CoV, coronavirus; COVID-19, coronavirus disease 2019; EtOH, ethanol; MERS-CoV, Middle East respiratory syndrome coronavirus; MLV, murine leukemia virus (MLV); MTT, 3-(4,5-dimethylthiazol-2-yl)-2,5-diphenyltetrazolium bromide; PCR, polymerase chain reaction; ROS, reactive oxygen species; RT-qPCR, real-time reverse transcription PCR; S, Spike; SARS-CoV, severe acute respiratory syndrome coronavirus; SARS-CoV-2, severe acute respiratory syndrome coronavirus 2; Tiron, 4,5-dihydroxy-1,3-benzene disulfonic acid; TTM, tetrathiomolybdate; VSV, vesicular stomatitis virus.

