## Supplemental Figs 1 to 4 for "Cellulosic copper nanoparticles and a hydrogen peroxide-based disinfectant protect Vero E6 cells against infection by viral pseudotyped particles expressing SARS-CoV-2, SARS-CoV or MERS-CoV Spike protein"

Table of contents

**Fig. S1**. *Effect of Saber™ and CMC-Cu on Vero E6 cell viability.* page 2

**Fig. S2**. *CMC-Cu inhibits SARS-CoV-2-S-pseudotyped virions.* page 3

**Fig. S3**. *Saber™ inhibits SARS-CoV-2-S-pseudotyped virions.* Page 4

**Fig. S4**. *The combined use of Saber™ and CMC-Cu inhibits SARS-CoV-2-S-pseudotyped virions.* Page 5

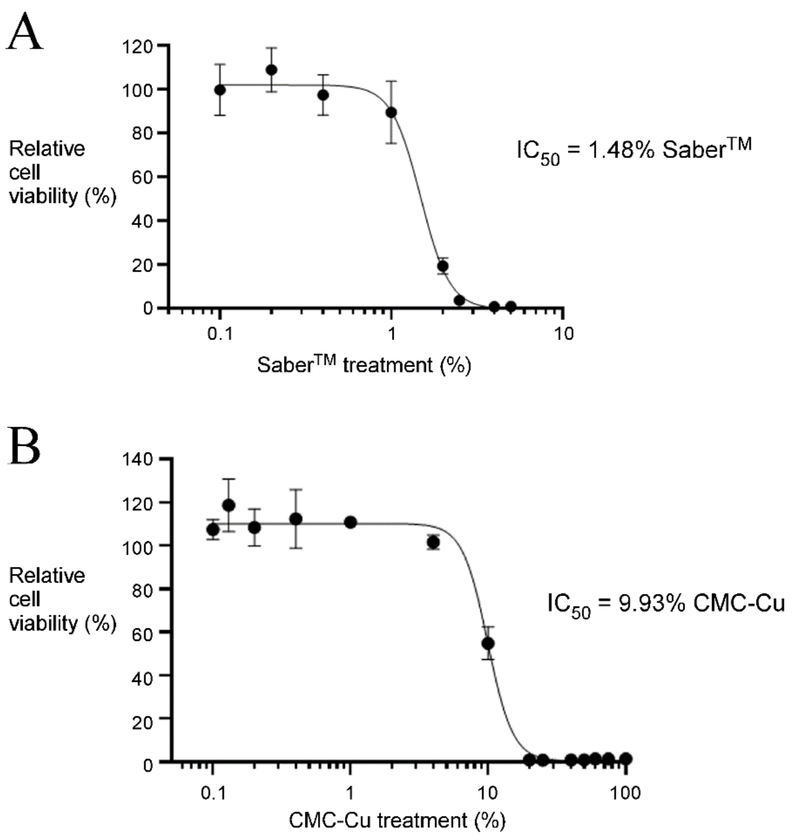


Figure S1
Brault *et al.*

**Fig. S1**. *Effect of Saber™ and CMC-Cu on Vero E6 cell viability. A – B,* Vero E6 cells were seeded in triplicate and treated with the indicated concentrations of Saber™ (*panel A*) or CMC-Cu (*panel B*) for 30 s. After gel filtration in the case of Saber™ and membrane filtration in the case of CMC-Cu, cells were further cultured for 72 h followed by measurements of cell viability by MTT assays.

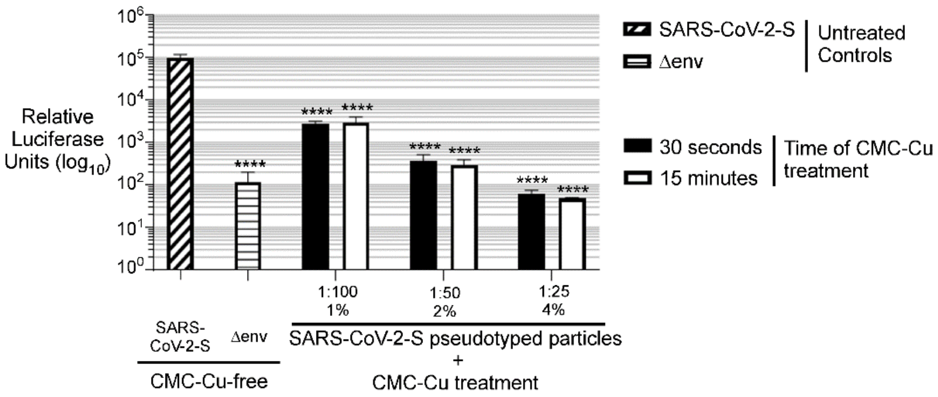


Figure S2
Brault *et al.*

**Fig. S2**. *CMC-Cu inhibits SARS-CoV-2-S-pseudotyped virions*. SARS-CoV-2-S-pseudotyped particles were exposed to CMC-Cu at concentrations of 1:100, 1:50, and 1:25 over a period of 30 s or 15 min prior to Vero E6 cell infection. As controls, Vero E6 cells were infected with untreated S-pseudovirions (CMC-Cu-free, positive control) or pseudovirions lacking a cell-surface viral envelope glycoprotein (Δenv, negative control). Luciferase assays were carried out 72 h post infection. Results are average relative luciferase units ($log_{10}$) of a minimum of five independent experiments performed in biological triplicate. Error bars indicate standard deviation (± SD; error bars). The asterisks correspond to $p < 0.0001$ (****) (one-way ANOVA with Dunnett's multiple comparisons test against the CMC-Cu-free SARS-CoV-2-S-pseudotyped particles).

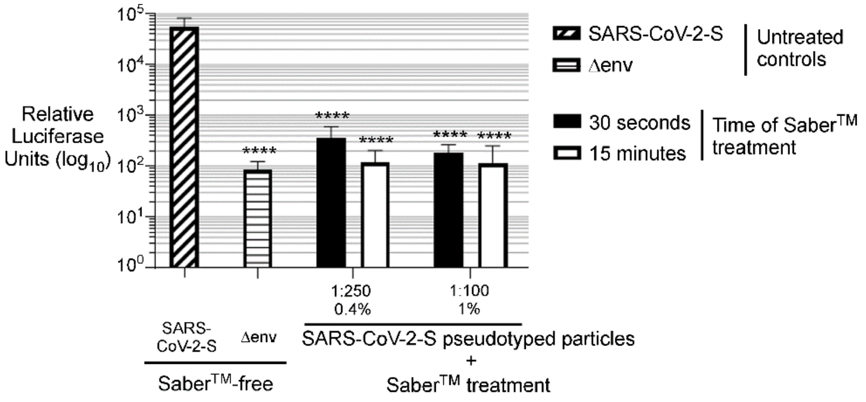


Figure S3

Brault *et al.*

**Fig. S3**. *Saber™ inhibits SARS-CoV-2-S-pseudotyped virions*. SARS-CoV-2-S-pseudotyped particles were incubated in the presence of Saber™ prepared in a specific volume of a dilute solution 1:250 (0.4%) and 1:100 (1%) from a concentrated stock solution. S-pseudovirions that were treated with Saber™ over a period of time of 30 s or 15 min were used to infect Vero E6 cells. As controls, Vero E6 cells were infected with untreated S-pseudoviral particles (Saber™-free, positive control) or pseudovirions lacking a cell-surface viral envelope glycoprotein (Δenv, negative control). Luciferase activity was measured 72 h post infection. Results are average relative luciferase units ($log_{10}$) of a minimum of five independent experiments performed in biological triplicate. Error bars indicate standard deviation (± SD; error bars). The asterisks correspond to $p < 0.0001$ (****) (one-way ANOVA with Dunnett's multiple comparisons test against the Saber™-free SARS-CoV-2-S-pseudotyped particles).

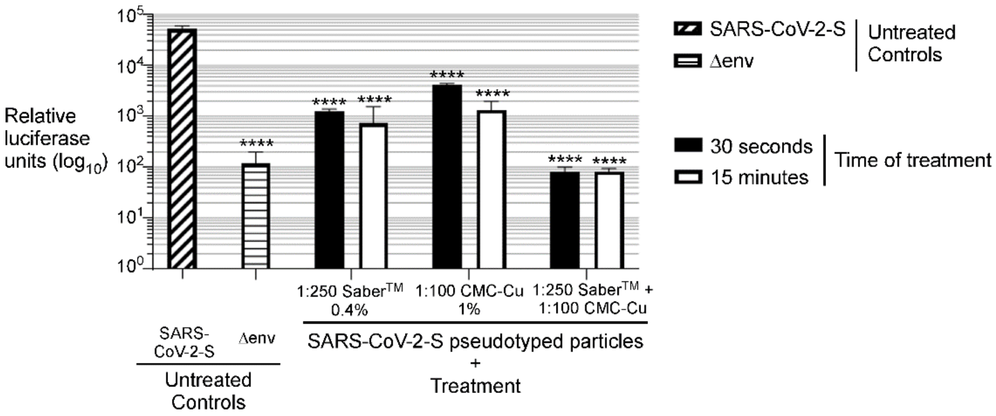


Figure S4
Brault *et al.*

**Fig. S4**. *The combined use of Saber™ and CMC-Cu inhibits SARS-CoV-2-S-pseudotyped virions*. SARS-CoV-2-S-pseudotyped virions were treated with Saber™ (1:250), CMC-Cu (1:100) or Saber™ in combination with CMC-Cu for 30 s or 15 min prior to Vero E6 cell infection. As controls, Vero E6 cells were infected with untreated S-pseudoviral particles (Saber™- and CMC-Cu-free, positive control) or pseudovirions lacking a cell-surface viral envelope glycoprotein (Δenv, negative control). Luciferase activity was measured 72 h post infection. Results are average relative luciferase units ($log_{10}$) of a minimum of five independent experiments performed in biological triplicate. Error bars indicate standard deviation (± SD; error bars). The asterisks correspond to $p < 0.0001$ (****) (one-way ANOVA with Dunnett's multiple comparisons test against the untreated SARS-CoV-2-S-pseudotyped particles).
